# Memory Traces Reflect How They Were Last Accessed

**DOI:** 10.1101/2025.05.30.657135

**Authors:** Xiaoxi Qi, Marc N. Coutanche

**Affiliations:** Department of Psychology, University of Pittsburgh, Pittsburgh, PA, USA; Learning Research and Development Center, University of Pittsburgh, Pittsburgh, PA, USA

## Abstract

Episodic memories are known to change with each act of retrieval. We hypothesize that accessing semantic knowledge in different ways during episodic retrieval—from unique perceptual features to taxonomic and thematic context—shapes how those memories are subsequently represented during recognition. In this functional magnetic resonance imaging (fMRI) study, human participants learned novel word–image pairs and underwent re-exposure that required accessing semantic knowledge at one of three different levels: item, category, or theme. Participants then completed a recognition task. Although behavioral memory performance was matched across conditions, neural activity during recognition varied based on prior semantic access history. Recognition patterns could be classified according to prior semantic access in early visual cortex, ventral temporal cortex and the hippocampus for remembered concepts. A whole-brain searchlight analysis revealed bilateral clusters along the ventral visual stream where semantic access history was decodable. To further characterize how prior semantic access history shaped neural representations during recognition, we tested whether memories accessed at the same semantic level were expressed more similarly or distinctly in the brain. Category-level access increased similarity among items in early visual cortex, while item-level access led to more differentiated representations in the visual word form area. Together, these findings show that how we access semantic knowledge during episodic retrieval leaves measurable traces in subsequent neural representations, revealing how semantic-episodic interactions shape memory representations.

## Introduction

Reconsolidation theory suggests that episodic memories are updated each time they are retrieved, making them sensitive to modification by external factors over time (e.g., Hupbach et al., 2007). Examples of such factors include altering temporal orders (St Jacques & Schacter, 2013) or retrieving with an alternative perspective (e.g., to reduce negative emotional components of a traumatic memory; James et al., 2015). Retrieval practice can enhance future behavioral memory performance (the testing effect; Kang et al., 2007; Roediger & Butler, 2011; Rowland, 2014) by reinforcing or creating new retrieval routes to memory representations (Roediger et al., 2011). When episodic memories are retrieved, the neural patterns initially activated during encoding are reactivated (Martin, 2007), potentially with new information being incidentally incorporated and the trace being updated (Forcato et al., 2010; Hupbach et al., 2007; J. L. C. Lee, 2009). Episodic memory retrieval often involves accessing semantic knowledge about the remembered concepts. Here, we ask how different patterns of semantic access during retrieval change episodic memory traces.

While episodic memories often include context-specific details (e.g., time, place, and emotional associations), semantic memories are typically independent of context and represent generalized knowledge (Tanguay et al., 2023). However, semantic memory is organized at multiple levels of granularity, ranging from specific features to broader categories (Lambon Ralph, 2014). Each level of detail contributes to the broader conceptual representation, influencing how and when the concept is used in future memory tasks (Fliessbach et al., 2010; Kuhnke et al., 2023). For instance, one’s semantic memory of a pet might encompass its name, species, and more (e.g., ’Dory’ is a ’Blue Tang,’ which is a species of fish, which is an animal). Although we may rarely *explicitly* retrieve this information, each of these semantic levels is essential for successfully interacting with objects and concepts. For instance, ‘Dory’ is a distinct entity from another Blue Tang and may have different nutritional requirements from its clown fish tank-mate. Each semantic granularity thus activates distinct characteristics of the concept (color vs. species, etc.) and may involve distinct neural activation patterns (Yee & Thompson-Schill, 2016). Multiple neurocognitive models suggest that conceptual knowledge representation is distributed across brain, with some models employing a hub that integrates modality-specific information to form coherent semantic representations (Lambon Ralph, 2014; Rogers et al., 2004). Such information is then accessed through regions such as the left ventrolateral prefrontal cortex (vlPFC), which engages in controlled semantic retrieval (Snyder et al., 2011).

Is there any reason for thinking that how we retrieve a concept might affect its memory trace? Retrieving item-level information (e.g., ‘Dory is blue’) can benefit future recognition of items (Dory vs. Charlie) while increasing the chance of false positives to similar items (lures; Brackmann et al., 2016; Bridge & Paller, 2012; St. Jacques et al., 2013). Retrieving thematic information (pet-related) more readily primes thematically related items (‘Fido’) than does retrieving categorical information (‘fish’ priming ‘lobster’; Sachs et al., 2008). These findings suggest that selectively engaging with different types of semantic knowledge may influence subsequent memory uses of related concepts. However, the neural mechanisms by which semantic retrieval at different granularities influences memory updating remain unclear.

Here, we test the hypothesis that accessing semantic knowledge in different ways during episodic retrieval shapes how memory traces are subsequently represented during recognition. In this study, each concept is accessed in one of three ways: through item-specific features, at the taxonomic (category) level, or at the thematic level. These three levels are thought to reflect qualitatively distinct forms of semantic processing and representation, especially taxonomic versus thematic access, which are often seen as being supported by partially separate neural systems (Mirman et al., 2017). Human participants retrieved learned information through one of these access routes, and we identified changes to the memory trace using functional magnetic resonance imaging (fMRI) with multivariate pattern analysis (MVPA), which allows the probing of fine-grained distinctions in neural memory representations (Coutanche, 2013; Haxby et al., 2001). We hypothesized that accessing different types of semantic information during episodic retrieval would create distinct neural signatures that persist during subsequent recognition. Specifically, we predicted that regions involved in controlled semantic retrieval (left vlPFC; Wing et al., 2013) would show modulated activity depending on prior semantic access. Accessing category-level information was predicted to emphasize shared perceptual features during recognition, leading to increased similarity in perceptual regions such as early visual cortex (EVC) and VT (Bainbridge et al., 2021). This prediction follows from the nature of taxonomic relations, which are typically grounded in shared perceptual features and preferentially engage visual regions (Kalénine et al., 2009; Mirman et al., 2017). Accessing a concept at the category level may therefore emphasize these shared perceptual features and increase the similarity of memory representations in perceptual regions during subsequent recognition.

## Materials and methods

### Participants

Forty-one participants were recruited for the study. Because of the importance of participants learning introduced information, we employed performance-based exclusion criteria based on surpassing chance-level performance in each task, consistent with approaches used in recent memory studies (e.g., Biderman & Shohamy, 2021; Skalaban et al., 2022; Smithson et al., 2023). Participants were excluded if their accuracy fell below 55% in either the localizer or the re-exposure task (both had 50% chance level for individual trials). These thresholds ensured that participants had successfully learned the introduced information necessary for reliable ROI localization and meaningful experimental manipulation. Twenty participants were excluded for low behavioral accuracy (1 participant from localizer, 18 from re-exposure, and 1 from both), and one was removed due to excessive motion. Therefore, twenty participants (13 females; age: Mean (M) = 24.4, standard deviation (SD) = 6.1) were included for data analyses. This sample size was consistent with recent fMRI studies examining semantic and episodic memory representations, which report reliable decoding and representational effects with samples of approximately 18–25 participants (e.g., Bainbridge et al., 2021a; S.-H. Lee et al., 2019; Wing et al., 2013). Participants provided written informed consent and were compensated for their time. All participants, aged between 18 and 39 years old, were right-handed with normal or corrected-to-normal vision, and had no reports of psychiatric or neurological conditions. Participants met the standard MRI safety requirements of the neuroimaging center, including an absence of metal in the body and a low likelihood of pregnancy. All participants were native English speakers with no previous experience with German and/or Dutch languages. The study was approved by the university’s Institutional Review Board.

### Stimuli

Participants were presented with pairings of object images and Dutch words. All Dutch words were composed of 3-8 letters based on the normalized dataset from Tokowicz et al. (2002). All words were concrete nouns, had one English translation, and had low form-similarity between Dutch words and their English translation, as rated by 24 Dutch-English bilinguals (Tokowicz et al., 2002). The images included items from 12 themes (camping, circus, farming, garbage, garden, ocean, picnic, summer, trees, vampire, vet, winter) and 4 object categories (animal, food, furniture, manipulable objects). There were 192 images in total, including 3 foils for each item that were used in the final recognition test. All word-image pairs were presented twice across two runs for each of the encoding, re-exposure, and recognition tasks. All images were scaled to the same size (250 x 250 pixels) and centered on a white background.

### Experimental procedure

After consenting, participants completed out-of-scanner behavioral category training and foil exposure, followed by five tasks during an fMRI scan (Task sequence is displayed in Figure 1 for all tasks, Figure 2 for before scan tasks, and Figure 3 for in-scanner tasks). The trial order for in-scanner tasks was pseudorandomized in advance, following the optimal sequence determined by the jitter optimization program Optseq 2 (https://surfer.nmr.mgh.harvard.edu/optseq/). After the scan, participants completed an out-of-scanner cued recall task.

**Figure 1.**
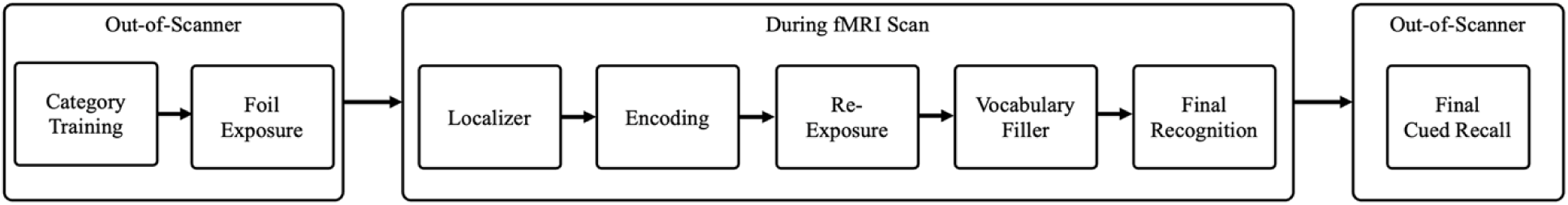
Task sequence acorss the study. Layout of the study procedure, including both out-of-scanner and in-scanner tasks. Participants completed category training and foil exposure before scanning, followed by a localizer, encoding, re-exposure, vocabulary filler, and final recognition task during a fMRI scan. A final cued recall task was completed after scanning.

**Figure 2.**
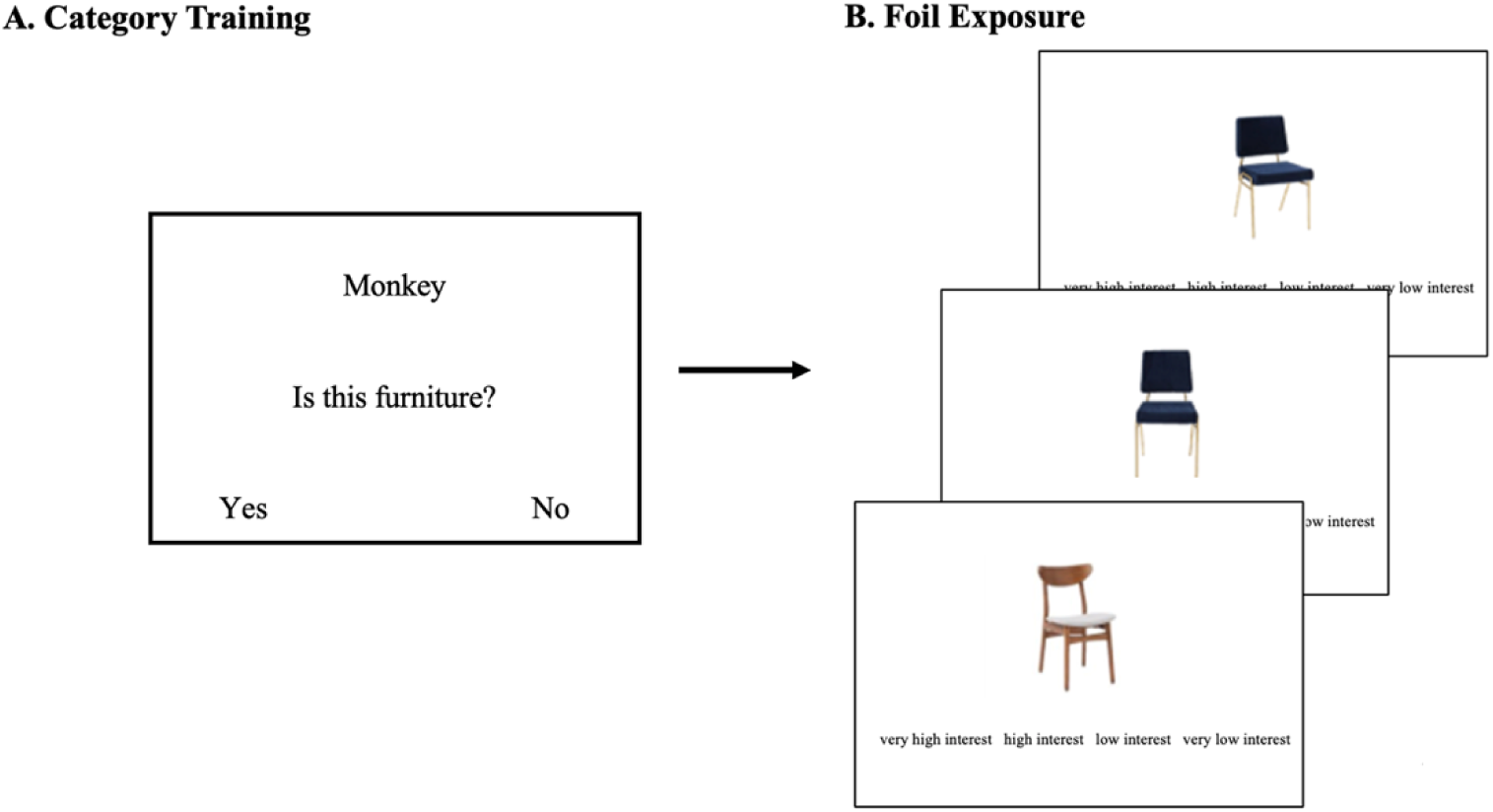
Tasks before scanning. (A) During category training, participants judged whether a word (e.g., “Monkey”) belonged to a target category (e.g., furniture), helping to familiarize them with object category definitions used in the study. (B) Foil exposure involved rating foil images on an interest scale. The example shows three foils for the target chair image, presented in the following order: correct exemplar with incorrect viewpoint, incorrect exemplar with correct viewpoint, and incorrect exemplar with incorrect viewpoint.

**Figure 3.**
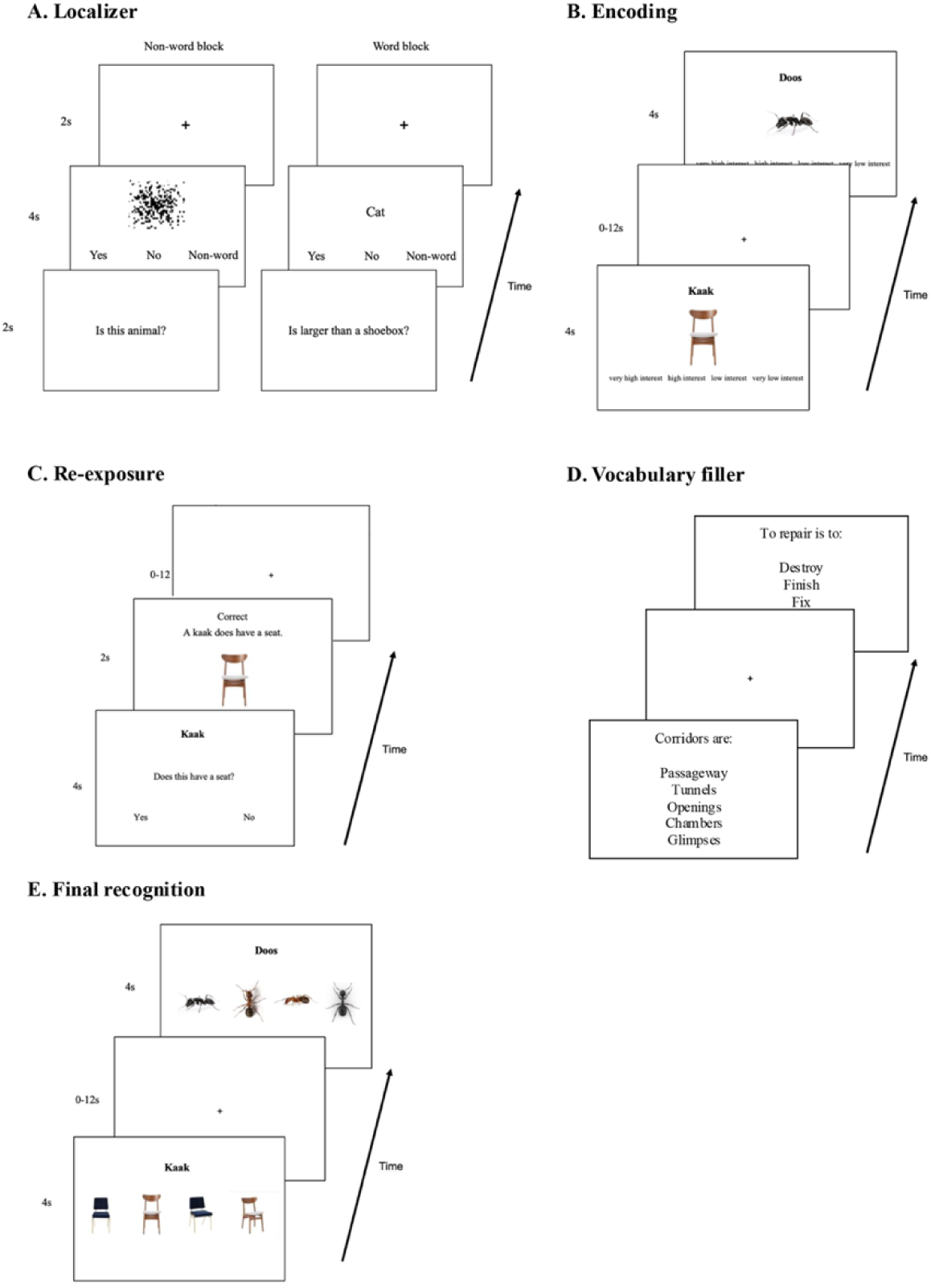
In-scanner task sequence. (A) Localizer task included alternating word and non-word blocks. Participants responded to a yes/no question at the beginning of each block (e.g., “Is this an animal?”), followed by real words or scrambled images. (B) During encoding, participants viewed word–image pairs and rated their interest. (C) In the re-exposure phase, participants answered a question targeting one of three semantic levels (e.g., “Does this have a seat?”) about the previously learned item. Corrective feedback followed each response. (D) The vocabulary filler task, shown during the anatomical scan, was selecting a synonym for a given word from four options. (E) During final recognition, participants matched each Dutch word to one of four images, which included the correct image and the same foil images previously seen during foil exposure.

#### Category training

We familiarized participants with the definitions of the object categories used in the study: animal, food, furniture, and manipulable objects. 12 known English words (3 for each category) were presented as examples, followed by a question targeting the category level of each word (e.g., “Is this food?”). Participants received feedback after each question (“Onion is a food. Foods can be consumed”). Words presented in this task did not overlap with items later paired with Dutch words.

#### Foil exposure

Participants were exposed to images of three future-foils for each of the items that would be paired with Dutch words. All foils appeared again in the final recognition. We included this foil exposure phase because previous studies indicated that initially exposing participants to foils would minimize potential novelty signals that would otherwise be elicited by foils during the final recognition task (Wing et al., 2013).

#### Localizer

We identified the visual word form area (VWFA) for each participant with a 10-minute localizer across 2 runs, each containing 6 blocks. There were two block types: word and non-word. At the start of each block, participants saw a yes/no question targeting a specific semantic level (e.g., “Is this an animal?”). In word blocks, participants viewed a series of real words and responded “yes” or “no” based on the question shown at the beginning of the block. In non-word blocks, participants viewed scrambled images and pressed a third button to indicate a “non-word” response. Each block began with the question screen presented for 2 seconds, followed by 8 stimuli shown for 4 seconds each. Stimuli were separated by two seconds of fixation. This task was used to identify regions sensitive to semantic granularity and word processing. To avoid interference, the words and questions used in this localizer task did not overlap with the stimuli used in the main tasks.

#### Encoding

Participants were asked to learn 48 pairs of item images and novel names (Dutch words). To avoid accidental correlations between phonologies and semantic features, the Dutch words were not translations of the item’s true names. Pairing randomization was counterbalanced across participants. Participants viewed each word-image pairing on the center of the screen for 4 seconds, with a question about their interest, in order to maintain engagement with the task (Figure 3). The pairings appeared once during each of the two functional runs, and trials were separated by a 0-12 second jittered interstimulus interval (M = 2 seconds).

#### Re-exposure

Each previously learned Dutch word was presented for 2 seconds with a yes/no question about the item associated with the word. Testing questions evoked one of the three target levels of semantic granularity (Table 1). Each pairing appeared twice in 2 runs and was only tested through one semantic level (e.g., testing *chair-ede* through category-level might include: “Is ede furniture?”/“Is ede animal?”). We provided feedback after each trial (e.g., “Correct. An ede is furniture”/“Incorrect. An ede is not animal”), with the associated image shown on the screen as well, because corrective feedback can enhance subsequent memory performance (Kang et al., 2007). The feedback screen was presented for 2 seconds. If the participant did not respond within 2 seconds, the trial was considered “incorrect.”

**Table 1.**
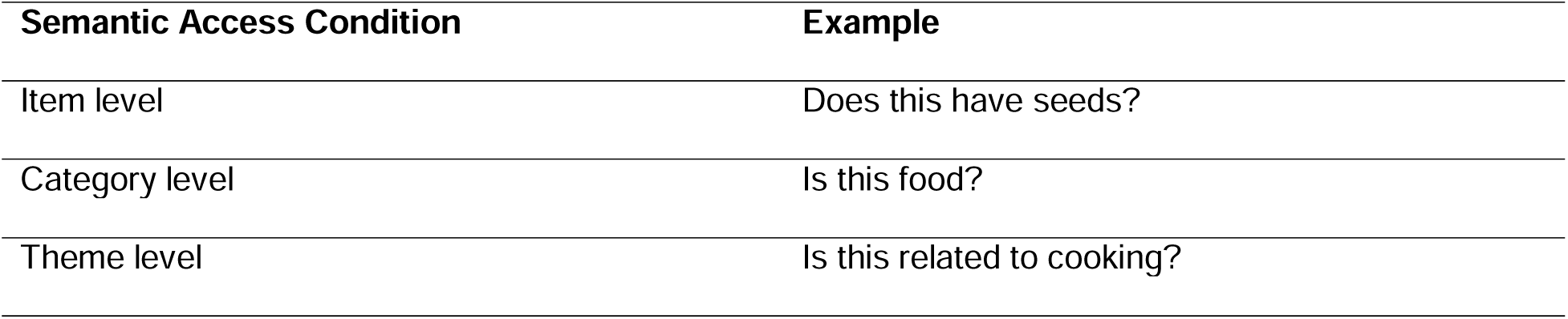
Example questions for semantic access conditions. This table provides example questions for each semantic access condition used in the re-exposure phase. The conditions represent semantic access at different levels: item, category, and theme. Each question was designed to prompt the participant to access a concept in semantic memory at the corresponding semantic level.

#### Vocabulary Filler

This task was administered during the anatomical scan. Participants were presented with a target word and asked to select which of four words most closely matched the target in meaning. This task was not relevant to the main analyses and was used to clear the working memory associated with the learned stimuli.

#### Recognition

Participants were asked to select the associated image for each Dutch word from four options, which included images of two similar exemplars and two viewpoints:

1. the correct exemplar from the correct viewpoint
2. the correct exemplar from an incorrect viewpoint
3. a similar lure exemplar from the correct viewpoint
4. a similar lure exemplar from an incorrect viewpoint

Each pairing was tested twice across two runs, with each trial lasting for 4 seconds.

#### Cued recall

After the scan, an out-of-scanner final cued recall task asked participants to type the associated Dutch word while viewing each previously paired image. Participants were instructed to give their best guess when spelling the words. Each trial ended when participants indicated that they had finished typing by pressing the “enter” key.

### Image acquisition

All scanning was performed on a Siemens 3-T Prisma MRI scanner equipped with a mirror device to perform fMRI stimuli presentation. Anatomical T1-weighted whole-brain MRI images were collected between re-exposure and final recognition phase (TR =1.540 s, TE =3.04s, voxel size = 1.0 x 1.0 x 1.0 mm). Data were collected over eight functional runs in total, with two runs per task. All functional runs used a multiband imaging sequence (multiband acceleration factor = 3) with whole-brain coverage (voxel size = 2.0 × 2.0 × 2.0 mm, TR = 2000 ms, TE = 30 ms). Each functional run included a 12-second (6 TR) buffer period after the task. For tasks after the localizer, a jittered interval between 0 and 12 seconds (mean = 2 seconds) was used to support estimation in the rapid event-related design, with the sequence optimized using Optseq 2 (https://surfer.nmr.mgh.harvard.edu/optseq/)

### Image preprocessing

Imaging data were processed using the Analysis of Functional NeuroImages (AFNI) software package (Cox, 1996). Structural scans were skull-stripped and standardized to the MNI standard space. Motion correction was applied to all functional images and registered to a mean functional volume. One participant was removed due to excessive motion. Functional data were smoothed using a kernel with full width at half maximum (FWHM) of 6mm for the univariate analyses. We only analyzed functional data recorded during recognition in this manuscript.

### Regions of interest

We defined eight regions of interest (ROIs) based on the semantic processing and word learning literature: VT, VWFA, anterior temporal lobe (ATL), left vlPFC, hippocampus, perirhinal cortex (PrC), medial parietal cortex (MPC), and early visual cortex (EVC) (Figure 4).

**Figure 4.**
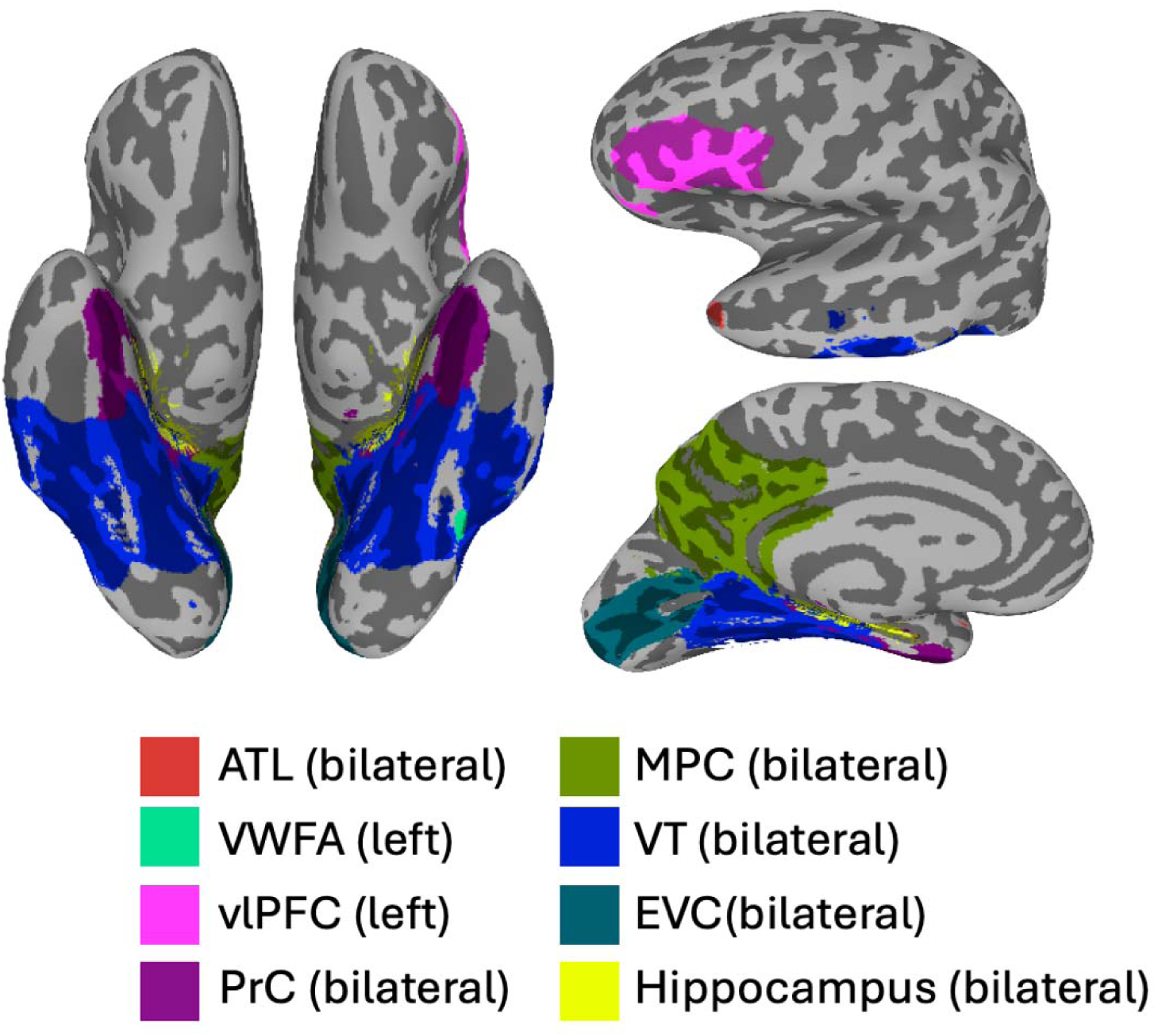
ROIs on a MNI template brain. Surface visualization of all regions of interest (ROIs) used in the study, shown on a standard MNI brain. ROIs include bilateral ATL, VT, EVC, MPC, PrC, hippocampus, and left-lateralized VWFA and vlPFC. Each ROI is color-coded and labeled below for reference.

The bilateral VT cortex codes complex visual stimuli and includes category-selective regions like the fusiform face and body-parts areas (FFAs and FBAs) (Haxby et al., 2011). To define VT, we used anatomical criteria based on prior literature: extending from 70 to 20 mm posterior to the anterior commissure in Talairach coordinates, incorporating the lingual, fusiform, parahippocampal, and inferior temporal gyri (Haxby et al., 2001). The VWFA is sensitive to word orthographic regularities, such as real words and pseudowords (Cohen et al., 2002; McCandliss et al., 2003), and was defined using a localizer task to identify regions near the left fusiform gyrus sensitive to word vs. non-word contrast, creating a 6 mm-radius sphere around the activity peak. The ATL functions as a semantic memory hub connecting multiple modalities (Lambon Ralph, 2014; Rice et al., 2015). We defined ATL using coordinates from prior studies, forming 6 mm-radius spheres at MNI coordinates (after conversion): 43, 13, -25; -42, 13, 23 (Coutanche & Thompson-Schill, 2015). The left vlPFC supports speech comprehension and semantic retrieval (Nozari & Novick, 2017) and was defined using Brodmann areas BA44, BA45, and BA47 from a Brodmann atlas (Mai, 2017). For the hippocampus and PrC, important in memory use (Davachi et al., 2003), and the MPC, associated with successful memory recall (Gilmore et al., 2015) and preferences for stimuli categories like people and places (Silson et al., 2019), we used Freesurfer atlas parcellations. The MPC specifically includes the precuneus, subparietal sulcus, and posterior cingulate gyrus. Early visual cortex has been modulated by top-down semantic and mnemonic processes (e.g., Ehrlich et al., 2024; Sergent et al., 2011) and we have previously found that EVC patterns can be shifted by learning perceptually-relevant dimensions (such as real-world size; Coutanche & Thompson-Schill, 2019). EVC was defined using Brodmann area BA17.

In our analyses, we selected the top 500 most active voxels from each large ROI (VT, MPC, PrC, hippocampus, and EVC). These voxels were selected by averaging voxel activity levels across all conditions (i.e., orthogonally) across two runs for pattern reactivation and pattern robustness analyses. For classification analyses, average activation in the training (but not testing) set was used to select 500 voxels for each fold.

### Behavioral analyses

Performance on the recognition task was measured as the proportion of “remembered” trials out of all recognition trials, with both recognition trials of each pairing contributing to the score. A “remembered” trial was characterized by the participant’s ability to correctly identify the object from the correct viewpoint. Recognition reaction times (RTs) were also recorded. Recognition accuracy and RTs were separately compared across semantic access conditions using one-way repeated-measures ANOVAs, with follow-up pairwise t-tests where the effect was significant. Performance on the cued recall task was measured on a continuous scale by calculating orthographic similarity scores between the word typed by the participant and the correct word paired with the images shown, using an online resource (van Orden, 1987).

Beyond overall recognition accuracy, we examined whether the type of error participants made depended on prior semantic access history. Each incorrect recognition response was classified into one of three error types based on the option selected: choosing the correct exemplar in the incorrect viewpoint (Type A), the incorrect exemplar in the correct viewpoint (Type B), or the incorrect exemplar in the incorrect viewpoint (Type C). Trials with no response were excluded. For each participant, we computed the proportion of each error type out of their total errors within each semantic access condition. These proportions were entered into a 3 (semantic access history: item, category, theme) × 3 (error type: A, B, C) rmANOVA, with participants as the random intercept. Follow-up pairwise comparisons between access conditions were conducted separately for each error type and corrected using the Benjamini-Hochberg FDR method.

Performance during re-exposure was scored in two ways because each pairing was re-exposed across two trials with corrective feedback after each. Both-trials accuracy was scored as the proportion of pairings answered correctly on both re-exposure trials. Second-trial accuracy was the proportion answered correctly on the second trial, which followed corrective feedback on the first trial. Re-exposure accuracy was compared across semantic access conditions using a one-way repeated-measures ANOVA for each scoring method, with follow-up pairwise t-tests where the effect was significant. We also examined whether final recognition accuracy depended on whether a pairing had been correctly accessed during re-exposure, see Supplementary Text S1 for details and results.

### Univariate analyses

A general linear model (GLM) was utilized to assess the BOLD signal associated with each type of semantic access history. Motion parameters were controlled by being included in the baseline model. No other nuisance regressors were included in the model. Stimuli were divided into 12 conditions based on object category and prior semantic access condition (e.g., animal retrieved via item-level question), and condition-level beta estimates were extracted during the final recognition phase.

#### Whole brain univariate activity

To identify regions sensitive to prior semantic access history across the whole brain, we first conducted a one-way rmANOVA testing for the main effect of semantic access history (item, category, theme), with participants as a random factor to account for individual differences in response patterns. This was followed by voxel-wise post-hoc paired t-tests comparing each pair of semantic access histories (item vs. category, item vs. theme, category vs. theme). Correction for multiple comparisons was applied using Monte-Carlo simulation with the 3dClustSim function in AFNI. A voxel-wise threshold of p < 0.005 and a cluster-wise alpha < 0.05 resulted in a corrected minimum cluster size of 72 voxels (edge-adjacent).

#### ROI-based univariate activity

We also assessed semantic access history effects within ROIs, where we tested for interactions between semantic access history and object category. A 4 (object category: animal, food, furniture, manipulable object) × 3 (semantic access history: item, category, theme) rmANOVA was conducted on the condition-level beta values from each ROI. Post-hoc paired t-tests were conducted in ROIs showing significant effects of semantic access history.

### Multivariate analyses

Functional data were not smoothed. Beta coefficients were calculated for each trial using the Least Squares-All (LSA, Mumford et al., 2012). Each voxel’s beta coefficients were z-scored across the time course of each run.

#### Recognition Pattern Classification

To test whether prior semantic access history (item, category, or theme) could be decoded from multivariate activity patterns during final recognition, we conducted classification analyses using a Gaussian Naive Bayes (GNB) classifier. We used a 2-fold procedure in which each run served as the training set in one fold and the testing set in the other, so the two trials of a pair were never in the same fold. The classifier performed a three-way classification to distinguish whether each stimulus had previously been retrieved at the item, category, or theme level (chance = 33%). This analysis was first performed using all trials in the recognition task, regardless of memory accuracy. Because each concept was tested on two trials, a concept was counted as correctly recognized if either of its two trials was correct, and both of its trials were then included. We then conducted a second analysis selected only correctly recognized trials. To ensure balanced class sizes (number of trials in each semantic access history) in this analysis with accuracy considered, we downsampled number of trials to match the class with the fewest instances for each participant. Participants with fewer than four included trials in any class (e.g., 6 trials each for category- and theme-level, but only 3 for item-level) were excluded from the analysis, resulting in a sample of 17 participants for this particular analysis. These analyses were performed both across the whole brain using a searchlight approach and within predefined regions of interest (ROIs).

##### Whole brain searchlight classification

was performed on local multivariate patterns extracted from 6-mm radius spherical searchlights centered on each voxel across the whole brain. Classification accuracy was assigned to the center voxel of each sphere, resulting in a whole-brain accuracy map for each subject. These maps were entered into a one-sample t-test, testing whether classification accuracy at each voxel exceeded chance level. Cluster-level correction for multiple comparisons was performed using the ClustSim option in 3dttest++, applying a voxel-wise threshold of p < 0.005 and a cluster-wise alpha of 0.05. This resulted in a minimum cluster size threshold of 119 voxels (edges adjacent) for corrected significance.

##### ROI-based classification

was performed within predefined ROIs. For larger ROIs (VT, PrC, MPC, hippocampus), we selected the top 500 most responsive voxels from the training set in each cross-validation fold to ensure consistent voxel sets between training and testing. For smaller ROIs, all voxels were used. Statistical significance was assessed using a permutation test. The testing labels were randomly shuffled 1000 times to generate a distribution of classification accuracies representing true chance performance. The classification accuracy observed with the correct labels was then compared to this distribution to assess significance. The p-value was calculated as the proportion of permutations that resulted in a value equal to or greater than the observed accuracy, then corrected for False Discovery Rate (FDR).

ROI-based classification was conducted in R (Version 4.4.2) using the e1071 package, and whole-brain searchlight classification was conducted in MATLAB (R2020a) using the Statistics and Machine Learning Toolbox. For each test trial, the classifier returned a posterior probability for each class. We retained the posterior probability of the true class as a trial-wise measure of classifier confidence, which was used in the following covariation analysis.

#### Covariation between Univariate Activity and Classifier Confidence Across ROIs

We then tested whether trial-wise univariate activity was related to the decodability of access history during recognition, using a form of informational connectivity (Anzellotti & Coutanche, 2018; Coutanche & Thompson-Schill, 2013). This analysis was conducted only with ROIs that either showed significant univariate effects of access history or significantly above-chance classification accuracies. Univariate activities reflected how strongly a region is engaged overall on a trial. Classifier confidence reflected how distinctly each type of access history is represented during recognition. For each recognition trial, we extracted the mean univariate response in a given region. Then, we extracted the posterior probability assigned to the true class from the GNB classifiers in selected ROIs as a trial-wise confidence value, concatenated across folds. For each participant, we computed the Spearman correlation between region-averaged univariate values and classifier confidence, separately for each of the region pairs, and Fisher-Z transformed the correlation coefficients before group-level analysis. Significance was assessed with a permutation test: we shuffled averaged univariate values within-run 1000 times while holding the classifier confidence values fixed to build a null distribution of group-averaged covariation. Because only three region pairs were tested (univariate: left vlPFC; multivariate: VT, EVC, and hippocampus), we report uncorrected p-values.

#### Covariation in Classifier Confidence Across ROI

As a follow-up to the ROI-based classification analysis, we examined whether regions that successfully decoded prior semantic access history showed coordinated trial-wise fluctuations in classifier confidence, also using an informational connectivity approach. This analysis was restricted to ROIs that achieved above-chance classification performance (VT, EVC, and hippocampus). See Text S2 for more details and results.

#### Recognition Pattern Consistency by Prior Semantic Access

While classification analyses established whether prior semantic access history could be decoded from recognition activity, they did not specify how this information is represented. To complement our classification analyses, we examined recognition pattern consistency to understand whether prior semantic access led to consistent updating of memory traces. Above-chance decoding could arise from various mechanisms (e.g., mean activation differences or changes in similarity), whereas pattern consistency specifically measures whether items with the same access history show systematic changes in their neural representation similarity, such as increased similarity among items sharing the same retrieval history, increased differentiation among those items, or both.

Recognition pattern consistency was calculated as “within-history” and “between-history” similarities of recognition-phase neural patterns. “Within-history” similarity was calculated by correlating recognition patterns of items that shared the same semantic access history from run 1 and run 2 (e.g., chair from recognition run 1 correlating to apple from recognition run 2, when both had been retrieved at the item-level during re-exposure). “Between-history” similarity was calculated by correlating recognition patterns of items with different semantic access histories across runs (e.g., chair from recognition run 1 correlating to bird from recognition run 2, when chair was accessed through item-level and bird through category-level during re-exposure).

The correlations of trials were always conducted across runs to avoid spurious correlations that might arise from within the same run (Mumford et al., 2014). Similarity scores were Fisher-Z transformed Spearman’s rank correlation coefficients. Because there were naturally more “between-history” than “within-history” correlations for each item, we permutated all “between-history” correlation coefficients (1000 times) and took a random sample (equal to the number of “within-history” correlations) in each iteration. The “within-history” and “between-history” similarities were averaged within each participant and compared against each other through group-level paired t tests. Because this involved 24 comparisons across ROIs and semantic access histories, p-values were corrected using the Benjamini-Hochberg FDR method. Effects that survived correction are reported as significant, but some uncorrected results are noted for descriptive purposes.

## Results

### Behavioral performance

Participants were able to learn the presented image-word pairings, as reflected by their final recognition (M = 0.50, SD = 0.12, chance= 0.25, t_(19)_ = 9.28, *p*<0.001) and cued recall (M = 0.45, SD = 0.17) performance. To ensure that any neural differences between the key conditions-of-interest are unlikely to result from systematic differences in memory performance across participants, we confirmed that group-level memory performance for trials in the various semantic access histories (item vs. category vs. theme questions) did not significantly differ in subsequent recognition (item: M = 0.52, SD = 0.13; category: M = 0.49, SD = 0.15; theme: M = 0.48, SD = 0.14; F_(2,38)_ = 0.98, p = 0.38) or cued recall (item: M = 0.43, SD = 0.20; category: M = 0.47, SD = 0.16; theme: M = 0.45, SD = 0.18; F_(2,38)_ = 1.23, p = 0.31). Recognition reaction times (RTs) did not differ across semantic access conditions (F(2,38) = 0.203, p = 0.817). RTs also did not differ between pairings that were correctly versus incorrectly accessed on the second re-exposure trial (t(19) = –0.53, p = 0.605).

We next asked whether the type of recognition error depended on prior semantic access history. The proportion of error types differed across access conditions, as indicated by a significant interaction between access history and error type (F(4, 68) = 2.79, p = 0.038). Neither main effect was significant. Follow-up pairwise comparisons between access conditions were conducted separately for each error type and FDR-corrected. No contrasts reached significance after correction (all p > 0.05). Descriptively, errors following item-level access tended to preserve exemplar identity, as participants were more likely to choose the correct exemplar but in the wrong viewpoint (Type A) after item-level access compared to theme-level access (t(19) = 2.36, p = 0.029, p_FDR_ = 0.087). Type C errors (incorrect exemplar, incorrect viewpoint) tended to be least frequent following item-level access and most frequent following theme-level access (item vs. theme: t(19) = –2.61, p = 0.017, p_FDR_ = 0.052; category vs. theme: t(19) = – 2.13, p = 0.047, p_FDR_ = 0.070).

We also examined performance during the re-exposure phase itself. Because each pairing was re-exposed across two trials, with corrective feedback after each trial, re-exposure performance can be characterized in two ways: accuracy on both trials of a pair, and accuracy on the second trial of a pair. Re-exposure accuracy varied across semantic access conditions when scored as both-trials-correct (F(2,38) = 3.61, p = 0.037), with item-level access showing higher accuracy than category-level access (t(19) = 2.55, p = 0.02); other pairwise comparisons were not significant (category vs theme: t(19) = –0.69, p = 0.50; item vs theme: t(19) = 1.81, p = 0.086). When scored as second-trial-correct, accuracy did not differ across conditions (F(2,38) = 0.82, p = 0.447). This suggests that participants learned from the corrective feedback, so that condition differences present on the first trial were resolved by the second trial.

### Recognition univariate activity

#### ROI-based univariate activity

Recognition task activity differed significantly based on how the trial’s concept was previously accessed in the left vlPFC (F(2,34) = 5.36, p = 0.01, p_FDR_= 0.117) with a significant interaction between access histories and object category (F(6,102) = 2.20, p = 0.049, p_FDR_= 0.221). However, none of these effects survived FDR correction for multiple comparisons across ROIs. No other ROIs demonstrated significant main effects or interaction effects (p >0.05).

#### Whole-brain univariate activity

To identify brain regions sensitive to semantic access history and examine the overall neural response pattern, we also conducted whole-brain univariate analyses investigating activation differences across semantic access conditions during recognition. We selected only successfully remembered trials to eliminate the confounding effect of behavioral accuracy. Clusters with significant differences based on semantic access history were located in the right inferior frontal cortex, bilateral ventral stream, bilateral middle occipital lobe, right superior frontal lobe, left lingual gyrus, and left precentral gyrus (Table 2). To examine which semantic access histories were contributing to each region’s result, post-hoc t-tests were conducted to compare the three levels (item, category, theme). Parts of the bilateral visual ventral stream, occipital lobe, and frontal cortex showed greater responses to one of the three forms of prior semantic access (Table 3; Figure 5).

**Figure 5.**
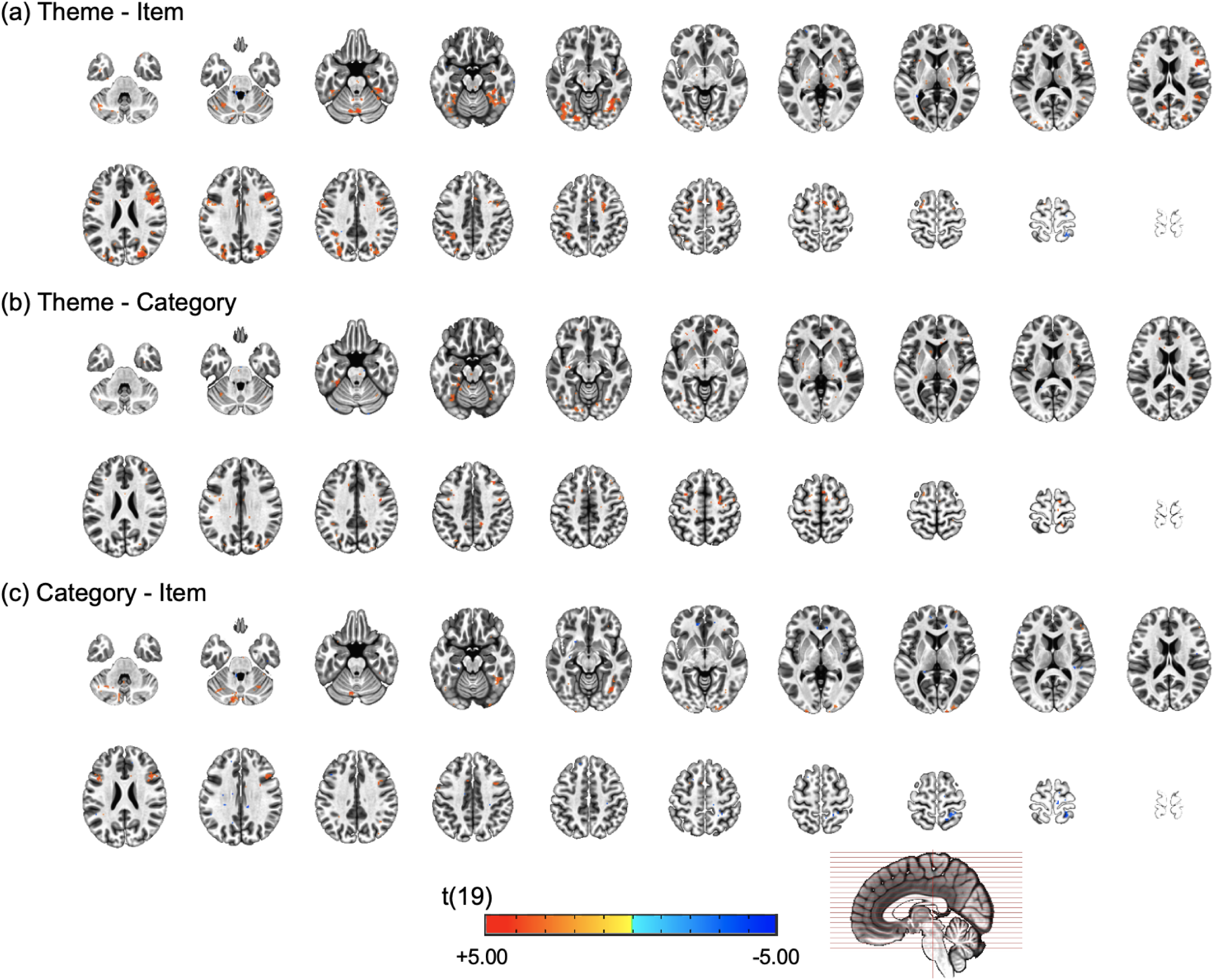
Whole-brain paired t-test contrasts between semantic access histories. Statistical maps of paired t-test results overlaid on an MNI template brain, shown for each pairwise contrast: (a) theme compared to item, (b) theme compared to category, and (c) category compared to item. Each contrast is displayed on 20 axial slices (every 6 mm) covering the whole brain. Color indicates the t-value at each voxel. All clusters survived a cluster-based correction for multiple comparisons at p < 0.005. The corresponding cluster coordinates and statistics are reported in Table 3.

**Table 2.** Significant clusters showed main effects of semantic access histories. Each cluster reflects a significant main effect of semantic access history in whole-brain analysis. Coordinates indicate the center-of-mass in MNI space. F-values correspond to the peak voxel within each cluster. M_Item_, M_Category_, and M_Theme_ show the mean beta estimate averaged across the cluster for each access condition, showing the direction of the effect across conditions. All clusters survived a cluster-based correction for multiple comparisons at p < 0.005.

| Region | #Voxels | x | y | z | Peak F | M<sub>Item</sub> | M<sub>Category</sub> | M<sub>Theme</sub> |
| --- | --- | --- | --- | --- | --- | --- | --- | --- |
| Right Inferior Frontal Gyrus | 626 | 45 | 18 | 26 | 6.96 | 0.13 | 0.28 | 0.34 |
| Left Fusiform Gyrus | 534 | -39 | -68 | -14 | 6.42 | 0.58 | 0.72 | 0.87 |
| Right Fusiform Gyrus | 511 | 37 | -56 | -16 | 7.95 | 0.52 | 0.67 | 0.78 |
| Right Inferior Occipital Gyrus | 304 | 30 | -77 | 28 | 7.63 | 0.36 | 0.47 | 0.57 |
| Right Superior Frontal Gyrus | 165 | 26 | -2 | 56 | 6.44 | 0.09 | 0.12 | 0.21 |
| Left Middle Occipital Gyrus | 158 | -29 | -78 | 34 | 7.3 | 0.34 | 0.45 | 0.57 |
| Left Lingual Gyrus | 155 | -18 | -85 | -10 | 8.94 | 0.50 | 0.61 | 0.79 |
| Left Cerebellum | 128 | -29 | -68 | -49 | 8.39 | 0.18 | 0.29 | 0.31 |
| Left Precentral Gyrus | 108 | -50 | 2 | 34 | 11.35 | 0.22 | 0.32 | 0.42 |
| Right Superior Parietal Lobule | 106 | 25 | -47 | 69 | 8.17 | 0.05 | -0.09 | -0.06 |
| Left Cerebellum | 100 | -6 | -73 | -28 | 9.95 | 0.22 | 0.33 | 0.36 |
| Left Inferior Parietal Lobule | 92 | -33 | -49 | 41 | 8.56 | 0.33 | 0.40 | 0.52 |

**Table 3.**
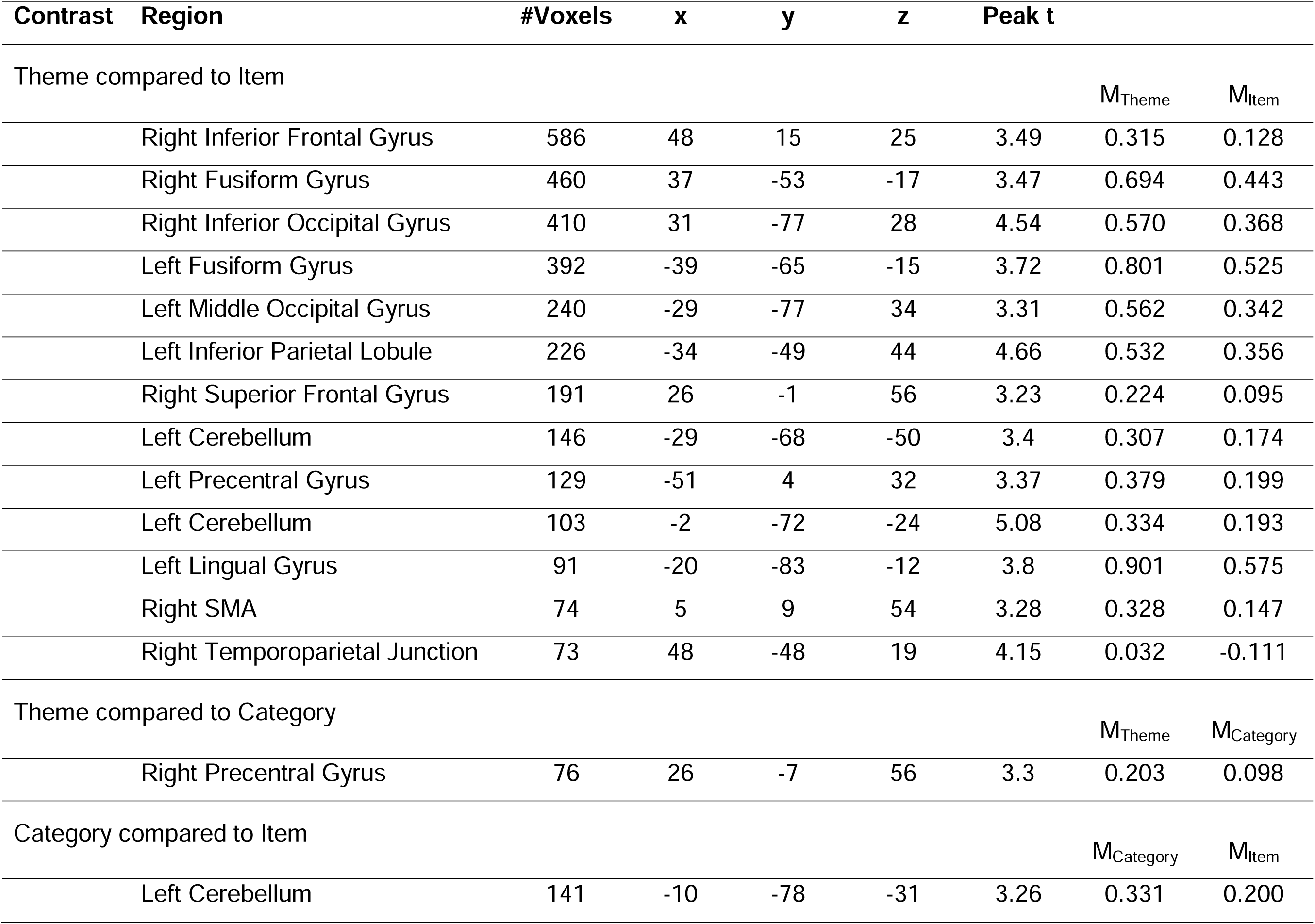
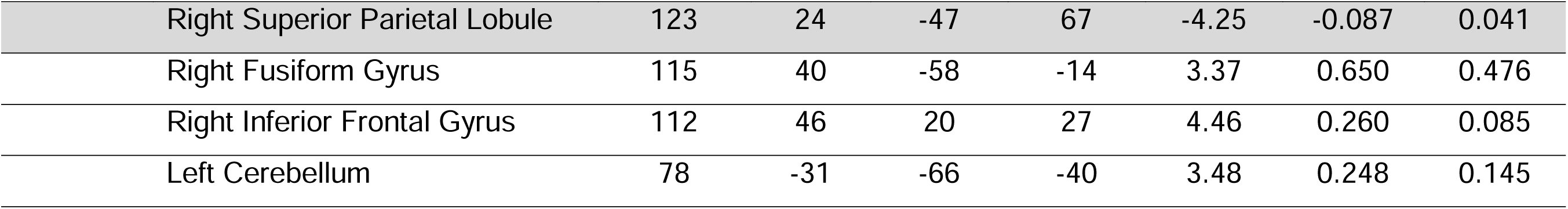
Significant cluster from paired t-tests on contrasts between semantic access histories. Each cluster reflects a significant difference between two semantic access conditions in whole-brain paired t-tests. Coordinates indicate the center-of-mass in MNI space, and t-values are reported at the peak voxel of each cluster. For each contrast, the two rightmost columns show the mean beta estimate averaged across the cluster for each of the two access conditions being compared, showing the direction of the difference. Negative t-values (i.e., effects in the opposite direction) are highlighted in gray. All clusters survived a cluster-based correction at p < 0.005.

### Recognition Pattern Classification

To test whether semantic access history creates distinguishable neural signatures that persist during subsequent recognition, we used multivariate pattern classification to decode prior semantic access conditions from recognition activity patterns.

#### Whole brain searchlight classification

To identify brain regions where recognition activity reflected semantic access history, we first conducted a whole-brain searchlight analysis that classified recognition trials based on their prior semantic access history (item, category, theme). The analysis revealed three clusters where classification accuracy exceeded chance level (33%) (Table 4; Figure 6): bilaterally in occipital cortex and in right lateral occipitotemporal cortex. These clusters included portions of primary visual cortex and extended anteriorly into fusiform gyrus in the right hemisphere. Mean classification accuracy across voxels within each cluster was 0.37 (SDs = 0.006–0.007), with peak searchlight accuracies ranging from 0.38 to 0.39. Reducing trials to correctly recognized did not identify clusters, likely because of the reduced power.

**Figure 6.**
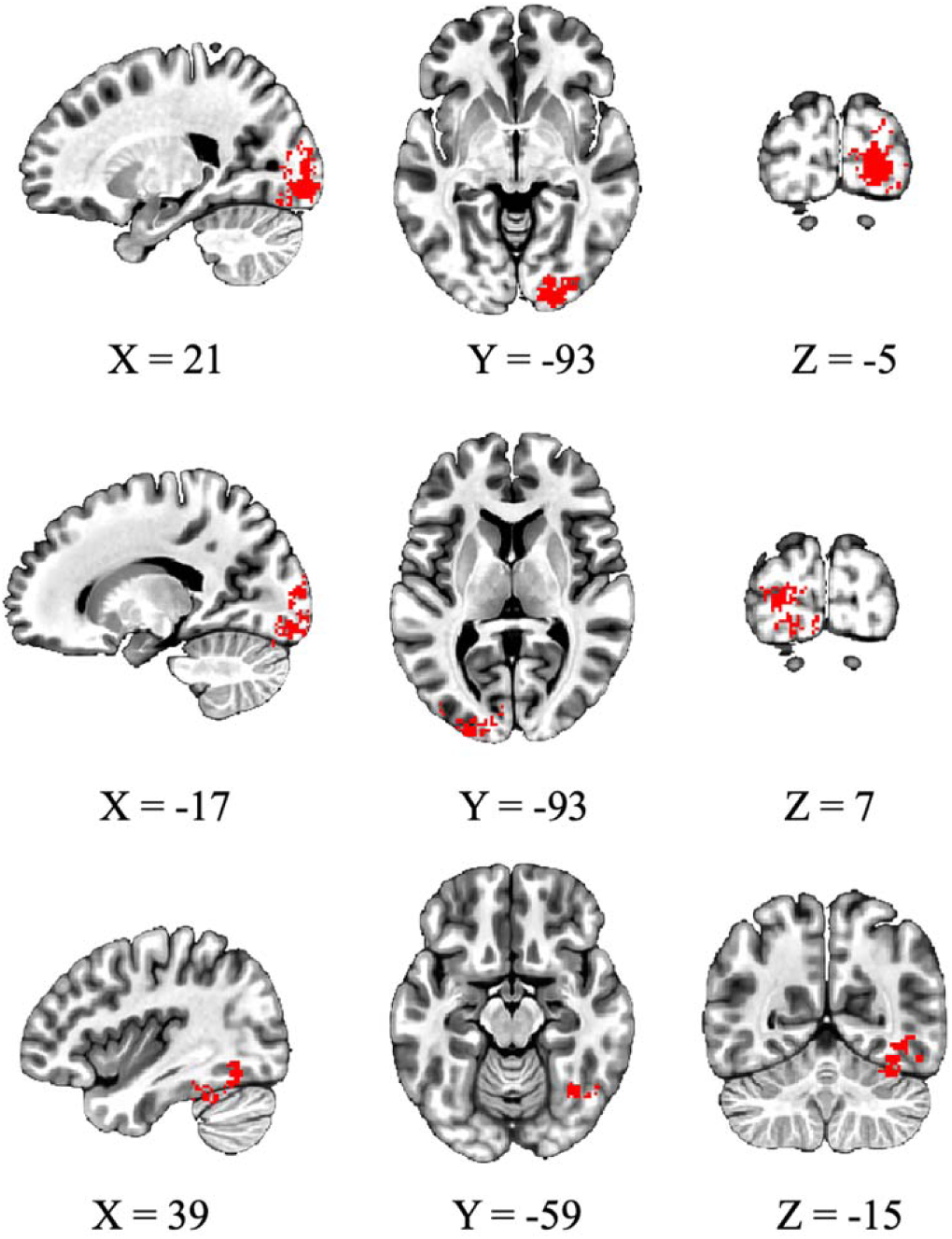
Significant clusters from whole-brain searchlight classification analysis. Example slices display brain regions where multivoxel pattern classification of prior semantic access history (item, category, or theme) during final recognition exceeded chance level (33%). Significant clusters are overlaid in red on an MNI template brain in LPI coordinate space. Each row presents a different cluster, with sagittal, axial, and coronal views shown for each. Coordinates were manually selected to optimize visibility of cluster locations.

**Table 4.**
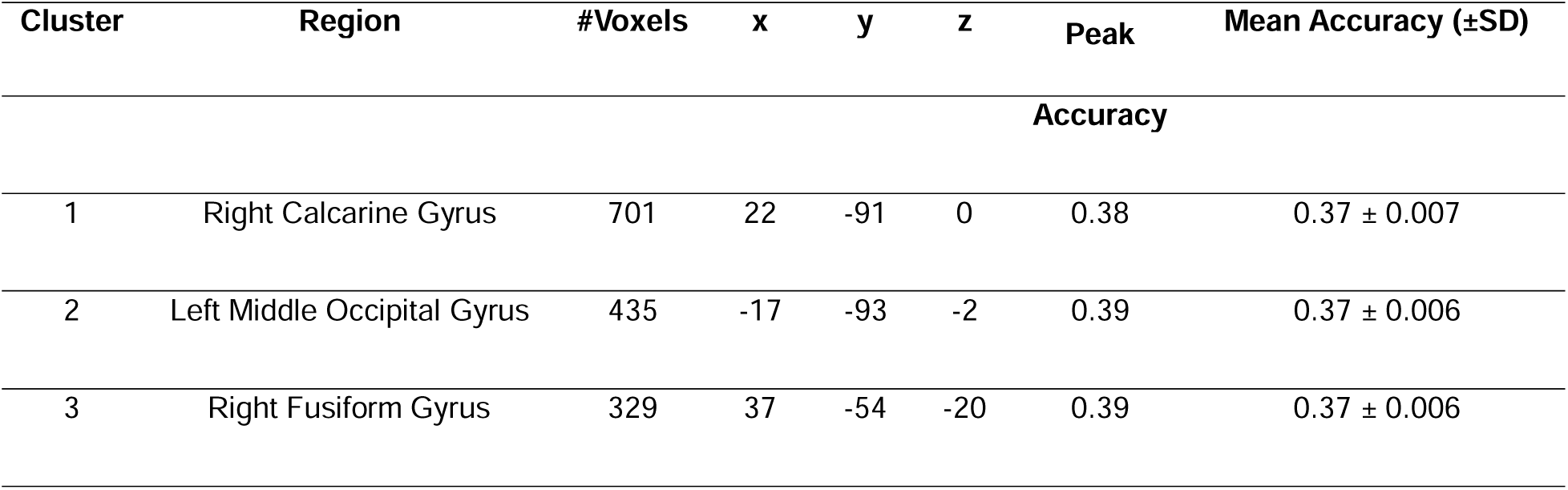
Significant clusters from whole-brain searchlight classification analysis. Clusters reflect regions where classification accuracy for prior semantic access history (item, category, or theme) during final recognition exceeded chance level (33%). Coordinates indicate the center-of-mass in MNI space. Peak and mean classification accuracies are reported for each cluster.

#### ROI based classification

Gaussian Naïve Bayes (GNB) classifiers were trained and tested on recognition neural patterns to decode semantic access history (item vs. category vs. theme levels; 33% chance, statistical significance calculated using permutation testing). Applied to the full set of trials, semantic access history could be decoded in VT (M = 0.35, SD = 0.05, p = 0.022, p_FDR_ = 0.124), EVC (M = 0.38, SD = 0.09, p = 0.009, p_FDR_ = 0.124), and was marginally significant in VWFA (M = 0.35, SD = 0.06, p = 0.08, p_FDR_ = 0.200) (Figure 7A), though these effects did not survive FDR correction for multiple comparisons across ROIs.

**Figure 7.**
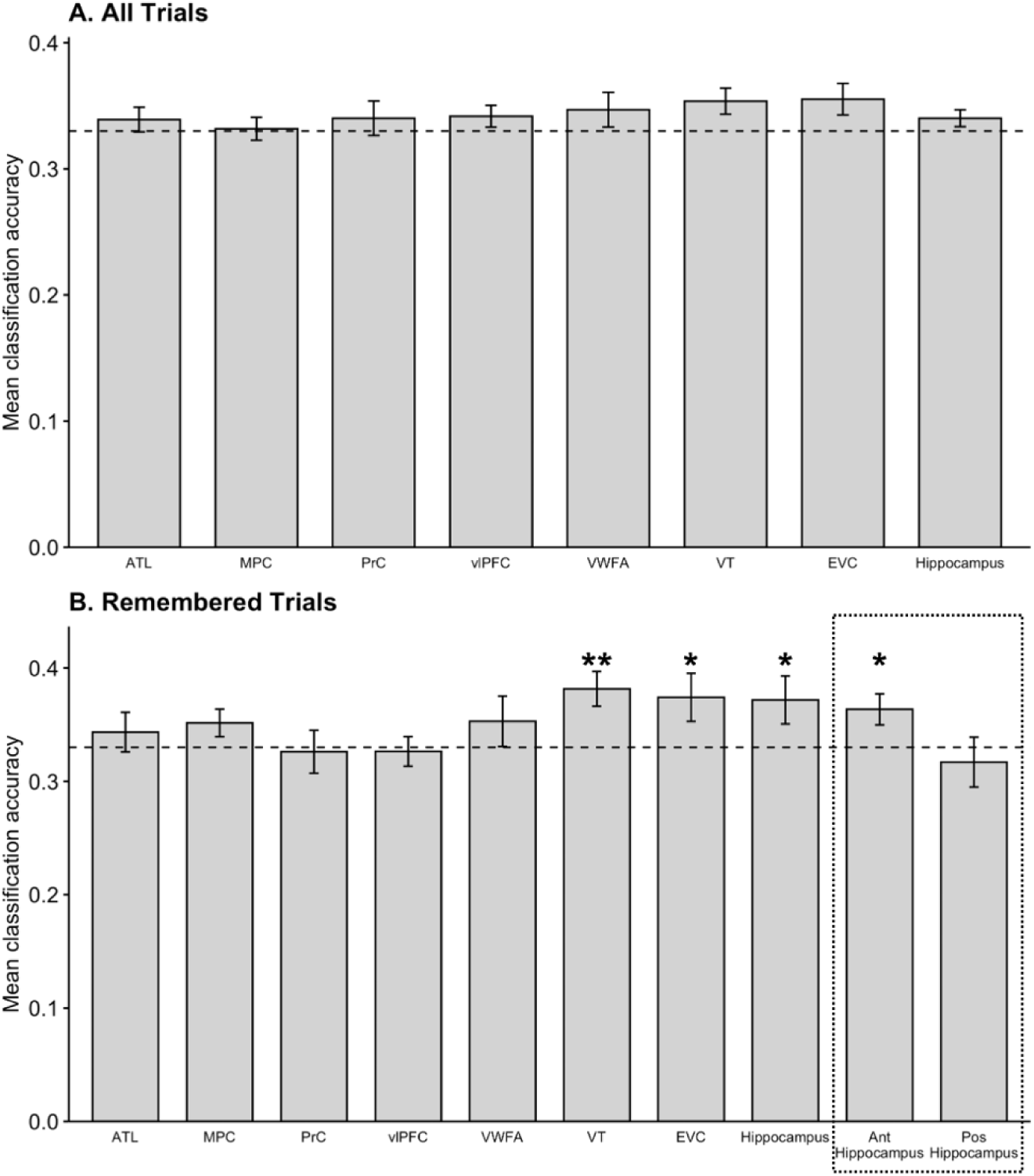
Three-way classification of semantic access history in recognition data. Mean classification accuracy for decoding prior semantic access history (item, category, theme) from multivoxel patterns during final recognition. Panel A includes all trials and panel B includes only correctly remembered trials. Bars represent accuracy for each ROI, and error bars indicate standard error of the mean. The dashed line represents chance level (33%). The dotted box groups the anterior and posterior hippocampus bars, indicating that these are subdivisions of the adjacent hippocampus ROI rather than separate regions. Significance values reflect group-level permutation tests: †*p* < .10, * *p* < .05, \*\**p* < .01, \*\*\**p* < .001.

Restricting the analysis to only successfully remembered trials, VT (M = 0.38, SD = 0.06, p<0.001, p_FDR_= 0.011), EVC (M = 0.36, SD = 0.06, p = 0.027, p_FDR_= 0.033), and the hippocampus (M = 0.37, SD = 0.09, p = 0.008, p_FDR_= 0.033) contained enough information to decode semantic access history (Figure 7B). To further investigate the hippocampus, the ROI was separated into anterior and posterior sections. The anterior hippocampus showed significant classification accuracy (M = 0.36, SD = 0.06, p = 0.036), while the posterior section did not show significant classification performance (M = 0.32, SD = 0.09, p = 0.859). The anterior sub-region had significantly greater decoding performance than the posterior sub-region (M = 0.05, SD = 0.10, p = 0.022) (Figure 7B).

### Covariation between Univariate Activity and Classifier Confidence Across ROIs

We tested whether trial-wise activity in the left vlPFC was related to classifier confidence in the hippocampus, EVC, and VT during recognition. Univariate activity in vlPFC was positively related to classifier confidence in the hippocampus (mean z = 0.08, p = 0.049). It was not related to classifier confidence in EVC (mean z = –0.007, p = 0.87) or VT (mean z = 0.004, p = 0.91). In other words, on trials with greater vlPFC engagement, access history was more decodable in the hippocampus, but not in the two perceptual regions.

### Recognition Pattern Consistency by Prior Semantic Access

We then examined whether prior semantic access led to consistent updating of memory traces, measured as increased similarity or differentiation among recognition-phase neural patterns for items sharing the same retrieval history. After FDR correction, EVC showed significantly greater similarity among items after category-level access compared to item- or theme-level access (t_(19)_ = 3.74, p= 0.001, p_FDR_ =0.017, Figure 8). This effect reflected increased similarity among items that shared a category-level semantic access history relative to items with different histories.

**Figure 8.**
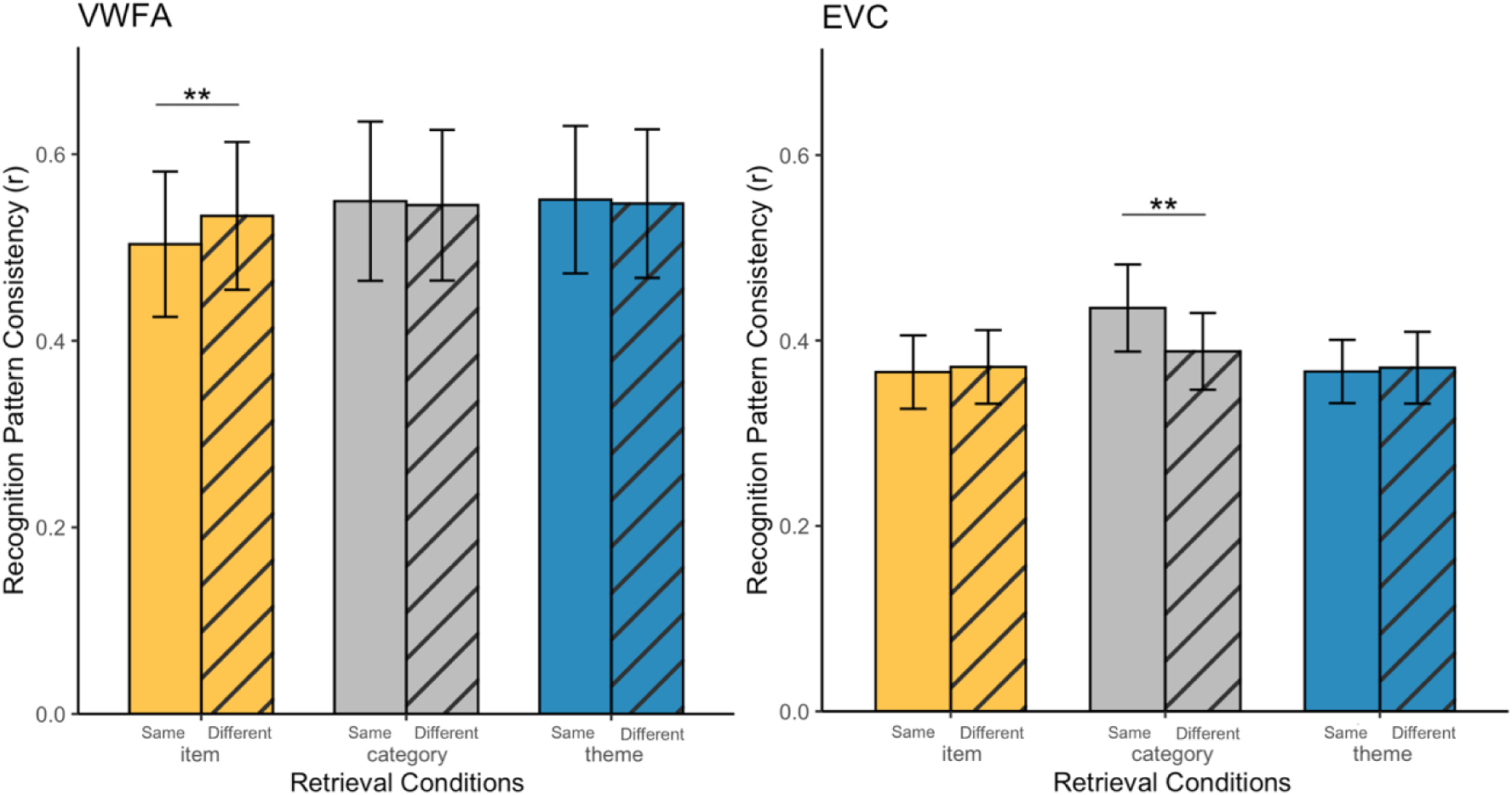
Significant recognition pattern consistency by prior semantic access results. Recognition neural patterns were compared for items previously tested through the same (within) versus different (between) semantic access conditions during re-exposure. Recognition pattern consistency was quantified as the difference in similarity between within-history and between-history item pairs, using Spearman’s correlation of recognition patterns and Fisher z-transformation prior to analysis. Results are shown for VWFA (left) and EVC (right). Error bars indicate group-level standard error of the mean. \*\**p* < .01, \*\*\**p* < .001.

After item-level access, VWFA remained the only region with a significant effect after correction, showing greater distinctiveness (lower similarity) compared to other semantic access histories (t_(19)_ = – 4.11, p <0.001, p_FDR_ = 0.007, Figure 8). Other item-level effects observed in the PrC (t_(19)_ = –2.24, p = 0.037), vlPFC (t_(19)_ = –2.37, p = 0.028), ATL (t_(19)_ = –1.89, p = 0.073), and hippocampus (t_(19)_ = –1.82, p = 0.08) were also in the negative direction but none of them survived FDR correction. These effects reflected reduced similarity among items with the same semantic access history following item-level access. No significant theme-level effects were observed in any region (p > 0.1, full results are provided in Table S1 and Figure S1).

## Discussion

We report new data showing that during memory recognition, neural representations carry an echo of how they were last accessed in semantic memory. By manipulating how concepts were accessed during an earlier phase and measuring neural activity during final recognition, we tested whether different types of semantic access leave distinct neural signatures. Notably, this manipulation reshaped neural representations during recognition while overall behavioral memory performance stayed unchanged. Neural pattern differences were evident across multiple analyses. Whole-brain univariate analyses revealed widespread effects of semantic access history across bilateral ventral stream and frontal regions, with theme-level access particularly engaging distributed neural networks compared to item-level access. Within ROIs, recognition activity in the left vlPFC differed by how the item had been accessed, though this effect was modulated by object category and did not survive correction across ROIs. Classification analyses demonstrated that these effects create reliably decodable patterns, with a whole-brain searchlight analysis identifying clusters in visual cortex where recognition activity could distinguish semantic access history above chance levels. The partial overlap between univariate and multivariate results suggests that semantic access history influences neural representations through both overall activation and more fine-grained pattern changes.

We next asked whether these regions carrying access-history information were functionally linked during recognition. Trial-wise univariate activity in the left vlPFC covaried with classifier confidence in the hippocampus, but not in EVC or VT. The left vlPFC supports the controlled retrieval and selection of semantic information (Wagner et al., 2001), and this link suggests that engagement of this control region was associated with how distinctly the hippocampus represented memories of access history. Trial-wise classifier confidence covaried across EVC, VT, and the hippocampus, indicating that coordinated expression of access history also exists across perceptual, conceptual, and memory-related regions (reported in Text S2).

We also tested whether accessing memories at the same semantic level led to consistent updating in how memory traces are activated during later recognition. We found that category-level access led to increased similarity among items in EVC, indicating that accessing general category information can create more overlapping neural representations, even in early visual areas. Category-level access likely emphasizes taxonomic semantic associations (e.g., “Dory is a fish”), making categorical features (e.g., “has fins”, “lives in water”) more accessible during subsequent recognition (Clarke & Tyler, 2014; Martin, 2007). Prior studies have shown that semantic expectations can influence early visual cortex through feedback signals (Bainbridge et al., 2021; Kok et al., 2012), and category-level information can be decoded from EVC under certain task demands (Cichy et al., 2014). Our findings add to this by showing that accessing category-level semantic knowledge leads to convergent updating of memory traces, causing increased similarity across items during subsequent recognition, possibly due to shared visual-semantic associations being reinforced.

In contrast, we observed that memory traces reshaped through item-level access (“Is Dory blue?”) were more distinct from one another during subsequent recognition than those accessed at the category or theme level. These findings suggest that accessing a concept with a focus on item-specific details enhances representational distinctiveness, reducing the similarity between items that were previously accessed in similar ways. This increased differentiation aligns with the concept of pattern separation, which is a process by which similar inputs are transformed into distinct, non-overlapping representations to avoid interference and aid future memory (LaRocque et al., 2013). Item-specific access in our study likely emphasized the features unique to each stimulus, reducing overlap among representations and increasing discriminability. This differentiation was found between concepts that were accessed at the item level. This fits our interpretation: when concepts are sharpened toward their own distinguishing features, their representations are pushed further apart than when an item-level concept is compared with one accessed at the category or theme level. These effects were significant in the VWFA after FDR correction, and present at an uncorrected level in regions including the PrC, vlPFC, ATL, and hippocampus. VWFA is associated with reading and orthographic processing and representing learned visual-verbal associations (Bruett et al., 2020; McCandliss et al., 2003). The PrC, which supports high-resolution object discrimination (Bussey et al., 2006), may similarly benefit from retrieval that emphasizes fine-grained item-specific features. As mentioned above, the left vlPFC supports controlled retrieval and selection of relevant semantic information (Wagner et al., 2001). These regions may help highlight the unique aspects of a memory and suppress more general or overlapping features (e.g., features that apply to many fish, but not Dory specifically). Together, these results show that item-level semantic access history enhances representational uniqueness, reducing the similarity of items in later recognition patterns even when they share contextual or categorical associations.

Importantly, recognition pattern consistency is not intended to provide a complete explanation of classification performance. Classification analyses establish that prior semantic access history is decodable during recognition, but they do not specify the representational mechanism that gives rise to this information. We observed partial dissociation across regions: significant consistency effects in VWFA without reliable classification, and significant classification in VT and hippocampus without corresponding consistency effects, suggesting that different brain regions preserve semantic access history through distinct representational changes. In some regions, such as EVC, decoding may be supported by convergent updating across items, while in VWFA, semantic access history may be expressed through increased differentiation without producing stable and consistent patterns that generalize across trials. Theme-level access produced the broadest univariate engagement across ventral and frontal regions, but did not show reliable changes in representational similarity in any region. This suggests that thematic access influenced the overall level of neural activity during recognition more than the fine-grained similarity structure of memory representation. Together, these findings indicate that different types of prior semantic access shapes memory representations during recognition through multiple mechanisms specific to each region rather than a single uniform process, with evidence of coordination across regions as reflected in the covariation analysis.

Participants learned the word–image pairings successfully and performed above chance at recognition, but recognition accuracy and reaction time did not differ across access conditions. They also did not differ between concepts that were correctly versus incorrectly accessed during re-exposure (reported in Text S1). Because corrective feedback followed every re-exposure trial, participants could learn a pairing even after an incorrect response, so re-exposure accuracy did not predict later recognition. This matched performance, and our use of only remembered trials balanced across conditions in the classification analyses, confirmed that the neural effects reflected memory representation rather than systematic differences in memory strength. However, these representational changes were still behaviorally relevant: although error rates did not differ across access conditions, the type of errors did. Errors following item-level access tended to preserve exemplar identity (and involved different viewpoints), while errors following theme-level access more often involved selecting the wrong exemplar entirely. This indicates that item-level access strengthened identity-specific information, such that even unsuccessful recognition retained the exemplar’s identity. This behavioral pattern converges with the greater neural distinctiveness following item-level access in VWFA described above.

We acknowledge that our experimental design prevents full within-stimulus control across conditions, as each stimulus is necessarily assigned to only one semantic access condition per participant. This means it is not possible to directly compare how the same stimulus is represented under different semantic access histories within the same individual. Additionally, our recognition pattern consistency analysis focused on correctly recognized trials. Examining incorrectly recognized trials would be valuable for understanding whether semantic access history effects persist even when memory traces are insufficient to support correct recognition. Past studies have shown that dynamic changes in neural representations can occur even in the absence of accurate memory performance, suggesting that false or inaccurate memories may still reflect meaningful updates to underlying memory traces (Shao et al., 2022; Zhuang et al., 2022). Future studies with larger trial counts would be needed to determine whether semantic access history effects also persist in incorrectly recognized trials, which could reveal the full extent of memory trace updating induced by re-exposure manipulations.

Taken together, these results suggest that different brain regions contribute in distinct ways to update memory traces based on the history of semantic access. Perceptual regions like EVC are more sensitive to category-level access, showing increased similarity between items that share taxonomic features. These regions may support the reinstatement of both perceptual and semantic contents used in earlier retrieval (Bainbridge et al., 2021). In contrast, language and memory-related areas, including the VWFA, PrC, and vlPFC, are more responsive to item-level access, showing greater distinctiveness across memories. These areas may help maintain more detailed and separate memory traces by focusing on item-specific features. This distribution across brain regions supports the idea that the brain preserves the history of how a memory was previously accessed in multiple ways (Ritchey & Cooper, 2020). By showing how item-, category-, and theme-level access histories are reflected in different brain areas, our study highlights the flexible and content-specific ways in which the brain stores and retrieves memories.

## Acknowledgements

This work was supported by a grant from the National Science Foundation (1947685).

## Author’s Note

This is the authors’ final version of the manuscript, incorporating all revisions made during peer review. The article has been accepted for publication in the *Journal of Cognitive Neuroscience*.

## References

1. Anzellotti, S., & Coutanche, M. N. (2018). Beyond Functional Connectivity: Investigating Networks of Multivariate Representations. Trends in Cognitive Sciences, 22(3), 258–269. 10.1016/j.tics.2017.12.002

2. Bainbridge, W. A., Hall, E. H., & Baker, C. I. (2021). Distinct Representational Structure and Localization for Visual Encoding and Recall during Visual Imagery. Cerebral Cortex (New York, N.Y.: 1991), 31(4), 1898–1913. 10.1093/cercor/bhaa329

3. Biderman, N., & Shohamy, D. (2021). Memory and decision making interact to shape the value of unchosen options. Nature Communications, 12(1), 4648. 10.1038/s41467-021-24907-x

4. Brackmann, N., Otgaar, H., Sauerland, M., & Howe, M. L. (2016). The impact of testing on the formation of children’s and adults’ false memories. Applied Cognitive Psychology, 30(5), 785–794. 10.1002/acp.3254

5. Bridge, D. J., & Paller, K. A. (2012). Neural correlates of reactivation and retrieval-induced distortion. The Journal of Neuroscience: The Official Journal of the Society for Neuroscience, 32(35), 12144–12151. 10.1523/JNEUROSCI.1378-12.2012

6. Bruett, H., Calloway, R. C., Tokowicz, N., & Coutanche, M. N. (2020). Neural pattern similarity across concept exemplars predicts memory after a long delay. NeuroImage, 219, 117030. (WOS:000559780900016). 10.1016/j.neuroimage.2020.117030

7. Bussey, T. J., Saksida, L. M., & Murray, E. A. (2006). Perirhinal cortex and feature-ambiguous discriminations. Learning & Memory, 13(2), 103–105. 10.1101/lm.163606

8. Cichy, R. M., Pantazis, D., & Oliva, A. (2014). Resolving human object recognition in space and time. Nature Neuroscience, 17(3), 455–462. 10.1038/nn.3635

9. Clarke, A., & Tyler, L. K. (2014). Object-Specific Semantic Coding in Human Perirhinal Cortex. Journal of Neuroscience, 34(14), 4766–4775. 10.1523/JNEUROSCI.2828-13.2014

10. Cohen, L., Lehéricy, S., Chochon, F., Lemer, C., Rivaud, S., & Dehaene, S. (2002). Language-specific tuning of visual cortex? Functional properties of the Visual Word Form Area. Brain: A Journal of Neurology, 125(Pt 5), 1054–1069. 10.1093/brain/awf094

11. Coutanche, M. N. (2013). Distinguishing multi-voxel patterns and mean activation: Why, how, and what does it tell us? Cognitive, Affective & Behavioral Neuroscience, 13(3), 667–673. 10.3758/s13415-013-0186-2

12. Coutanche, M. N., & Thompson-Schill, S. L. (2013). Informational connectivity: Identifying synchronized discriminability of multi-voxel patterns across the brain. Frontiers in Human Neuroscience, 7. 10.3389/fnhum.2013.00015

13. Coutanche, M. N., & Thompson-Schill, S. L. (2015). Creating Concepts from Converging Features in Human Cortex. Cerebral Cortex (New York, N.Y.: 1991), 25(9), 2584–2593. 10.1093/cercor/bhu057

14. Coutanche, M. N., & Thompson-Schill, S. L. (2019). Neural activity in human visual cortex is transformed by learning real world size. NeuroImage, 186, 570–576. 10.1016/j.neuroimage.2018.11.039

15. Cox, R. W. (1996). AFNI: Software for analysis and visualization of functional magnetic resonance neuroimages. Computers and Biomedical Research, an International Journal, 29(3), 162–173. 10.1006/cbmr.1996.0014

16. Davachi, L., Mitchell, J. P., & Wagner, A. D. (2003). Multiple routes to memory: Distinct medial temporal lobe processes build item and source memories. Proceedings of the National Academy of Sciences of the United States of America, 100(4), 2157–2162. 10.1073/pnas.0337195100

17. Ehrlich, I., Ortiz-Tudela, J., Tan, Y. Y., Muckli, L., & Shing, Y. L. (2024). Mnemonic But Not Contextual Feedback Signals Defy Dedifferentiation in the Aging Early Visual Cortex. Journal of Neuroscience, 44(16). 10.1523/JNEUROSCI.0607-23.2023

18. Fliessbach, K., Buerger, C., Trautner, P., Elger, C. E., & Weber, B. (2010). Differential effects of semantic processing on memory encoding. Human Brain Mapping, 31(11), 1653–1664. (WOS:000283641100003). 10.1002/hbm.20969

19. Forcato, C., Rodríguez, M. L. C., Pedreira, M. E., & Maldonado, H. (2010). Reconsolidation in humans opens up declarative memory to the entrance of new information. Neurobiology of Learning and Memory, 93(1), 77–84. 10.1016/j.nlm.2009.08.006

20. Gilmore, A. W., Nelson, S. M., & McDermott, K. B. (2015). A parietal memory network revealed by multiple MRI methods. Trends in Cognitive Sciences, 19(9), 534–543. 10.1016/j.tics.2015.07.004

21. Haxby, J. V., Gobbini, M. I., Furey, M. L., Ishai, A., Schouten, J. L., & Pietrini, P. (2001). Distributed and overlapping representations of faces and objects in ventral temporal cortex. Science, 293(5539), Article 5539. 10.1126/science.1063736

22. Haxby, J. V., Guntupalli, J. S., Connolly, A. C., Halchenko, Y. O., Conroy, B. R., Gobbini, M. I., Hanke, M., & Ramadge, P. J. (2011). A common, high-dimensional model of the representational space in human ventral temporal cortex. Neuron, 72(2), 404–416. 10.1016/j.neuron.2011.08.026

23. Hupbach, A., Gomez, R., Hardt, O., & Nadel, L. (2007). Reconsolidation of episodic memories: A subtle reminder triggers integration of new information. Learning & Memory, 14(1–2), 47–53. 10.1101/lm.365707

24. James, E. L., Bonsall, M. B., Hoppitt, L., Tunbridge, E. M., Geddes, J. R., Milton, A. L., & Holmes, E. A. (2015). Computer Game Play Reduces Intrusive Memories of Experimental Trauma via Reconsolidation-Update Mechanisms. Psychological Science, 26(8), 1201–1215. 10.1177/0956797615583071

25. Kalénine, S., Peyrin, C., Pichat, C., Segebarth, C., Bonthoux, F., & Baciu, M. (2009). The sensory-motor specificity of taxonomic and thematic conceptual relations: A behavioral and fMRI study. NeuroImage, 44(3), 1152–1162. 10.1016/j.neuroimage.2008.09.043

26. Kang, S. H. K., McDermott, K. B., & Roediger III, H. L. (2007). Test format and corrective feedback modify the effect of testing on long-term retention. European Journal of Cognitive Psychology, 19(4–5), 528–558. 10.1080/09541440601056620

27. Kok, P., Jehee, J. F. M., & de Lange, F. P. (2012). Less Is More: Expectation Sharpens Representations in the Primary Visual Cortex. Neuron, 75(2), 265–270. 10.1016/j.neuron.2012.04.034

28. Kuhnke, P., Kiefer, M., & Hartwigsen, G. (2023). Conceptual representations in the default, control and attention networks are task-dependent and cross-modal. Brain and Language, 244, 105313. 10.1016/j.bandl.2023.105313

29. Lambon Ralph, M. A. (2014). Neurocognitive insights on conceptual knowledge and its breakdown. Philosophical Transactions of the Royal Society B: Biological Sciences, 369(1634), 20120392. 10.1098/rstb.2012.0392

30. LaRocque, K. F., Smith, M. E., Carr, V. A., Witthoft, N., Grill-Spector, K., & Wagner, A. D. (2013). Global Similarity and Pattern Separation in the Human Medial Temporal Lobe Predict Subsequent Memory. Journal of Neuroscience, 33(13), 5466–5474. 10.1523/JNEUROSCI.4293-12.2013

31. Lee, J. L. C. (2009). Reconsolidation: Maintaining memory relevance. Trends in Neurosciences, 32(8), 413–420. 10.1016/j.tins.2009.05.002

32. Lee, S.-H., Kravitz, D. J., & Baker, C. I. (2019). Differential Representations of Perceived and Retrieved Visual Information in Hippocampus and Cortex. Cerebral Cortex, 29(10), 4452–4461. 10.1093/cercor/bhy325

33. Mai, J. K. (with Majtanik, M.). (2017). Human brain in standard MNI space: Structure and function : a comprehensive pocket atlas. Academic Press, an imprint of Elsevier. https://shop.elsevier.com/books/human-brain-in-standard-mni-space/k-mai/978-0-12-811275-5

34. Martin, A. (2007). The representation of object concepts in the brain. Annual Review of Psychology, 58, 25–45. 10.1146/annurev.psych.57.102904.190143

35. McCandliss, B. D., Cohen, L., & Dehaene, S. (2003). The visual word form area: Expertise for reading in the fusiform gyrus. Trends in Cognitive Sciences, 7(7), 293–299. 10.1016/S1364-6613(03)00134-7

36. Mirman, D., Landrigan, J.-F., & Britt, A. E. (2017). Taxonomic and Thematic Semantic Systems. Psychological Bulletin, 143(5), 499–520. 10.1037/bul0000092

37. Mumford, J. A., Davis, T., & Poldrack, R. A. (2014). The impact of study design on pattern estimation for single-trial multivariate pattern analysis. NeuroImage, 103, 130–138. 10.1016/j.neuroimage.2014.09.026

38. Mumford, J. A., Turner, B. O., Ashby, F. G., & Poldrack, R. A. (2012). Deconvolving BOLD activation in event-related designs for multivoxel pattern classification analyses. NeuroImage, 59(3), 2636–2643. 10.1016/j.neuroimage.2011.08.076

39. Nozari, N., & Novick, J. (2017). Monitoring and Control in Language Production. Current Directions in Psychological Science, 26(5), 403–410. 10.1177/0963721417702419

40. Rice, G. E., Hoffman, P., & Lambon Ralph, M. A. (2015). Graded specialization within and between the anterior temporal lobes. Annals of the New York Academy of Sciences, 1359(1), 84–97. 10.1111/nyas.12951

41. Ritchey, M., & Cooper, R. A. (2020). Deconstructing the Posterior Medial Episodic Network. Trends in Cognitive Sciences, 24(6), 451–465. 10.1016/j.tics.2020.03.006

42. Roediger, H. L., & Butler, A. C. (2011). The critical role of retrieval practice in long-term retention. Trends in Cognitive Sciences, 15(1), 20–27. 10.1016/j.tics.2010.09.003

43. Rogers, T. T., Lambon Ralph, M. A., Garrard, P., Bozeat, S., McClelland, J. L., Hodges, J. R., & Patterson, K. (2004). Structure and Deterioration of Semantic Memory: A Neuropsychological and Computational Investigation. Psychological Review, 111(1), 205–235. 10.1037/0033-295X.111.1.205

44. Rowland, C. A. (2014). The effect of testing versus restudy on retention: A meta-analytic review of the testing effect. Psychological Bulletin, 140(6), 1432–1463. 10.1037/a0037559

45. Sachs, O., Weis, S., Krings, T., Huber, W., & Kircher, T. (2008). Categorical and thematic knowledge representation in the brain: Neural correlates of taxonomic and thematic conceptual relations. Neuropsychologia, 46(2), 409–418. 10.1016/j.neuropsychologia.2007.08.015

46. Sergent, C., Ruff, C. C., Barbot, A., Driver, J., & Rees, G. (2011). Top–Down Modulation of Human Early Visual Cortex after Stimulus Offset Supports Successful Postcued Report. Journal of Cognitive Neuroscience, 23(8), 1921–1934. 10.1162/jocn.2010.21553

47. Shao, X., Chen, C., Loftus, E. F., Xue, G., & Zhu, B. (2022). Dynamic changes in neural representations underlie the repetition effect on false memory. NeuroImage, 259, 119442. (WOS:000863467900008). 10.1016/j.neuroimage.2022.119442

48. Silson, E. H., Steel, A., Kidder, A., Gilmore, A. W., & Baker, C. I. (2019). Distinct subdivisions of human medial parietal cortex support recollection of people and places. eLife, 8, e47391. 10.7554/eLife.47391

49. Skalaban, L. J., Cohen, A. O., Conley, M. I., Lin, Q., Schwartz, G. N., Ruiz-Huidobro, N. A. M., Cannonier, T., Martinez, S. A., & Casey, B. J. (2022). Adolescent-specific memory effects: Evidence from working memory, immediate and long-term recognition memory performance in 8–30 yr olds. Learning & Memory, 29(8), 223–233. 10.1101/lm.053539.121

50. Smithson, C. J. R., Eichbaum, Q. G., & Gauthier, I. (2023). Object recognition ability predicts category learning with medical images. Cognitive Research: Principles and Implications, 8(1), 9. 10.1186/s41235-022-00456-9

51. Snyder, H. R., Banich, M. T., & Munakata, Y. (2011). Choosing Our Words: Retrieval and Selection Processes Recruit Shared Neural Substrates in Left Ventrolateral Prefrontal Cortex. Journal of Cognitive Neuroscience, 23(11), 3470–3482. 10.1162/jocn_a_00023

52. St. Jacques, P. L., Olm, C., & Schacter, D. L. (2013). Neural mechanisms of reactivation-induced updating that enhance and distort memory. Proceedings of the National Academy of Sciences, 110(49), 19671–19678. 10.1073/pnas.1319630110

53. St Jacques, P. L., & Schacter, D. L. (2013). Modifying memory: Selectively enhancing and updating personal memories for a museum tour by reactivating them. Psychological Science, 24(4), 537–543. 10.1177/0956797612457377

54. Tanguay, A. F., Palombo, D. J., Love, B., Glikstein, R., Davidson, P. S., & Renoult, L. (2023). The shared and unique neural correlates of personal semantic, general semantic, and episodic memory. eLife, 12, e83645. 10.7554/eLife.83645

55. Tokowicz, N., Kroll, J. F., de Groot, A. M. B., & van Hell, J. G. (2002). Number-of-translation norms for Dutch—English translation pairs: A new tool for examining language production. Behavior Research Methods, Instruments, & Computers, 34(3), 435–451. 10.3758/BF03195472

56. van Orden, G. C. (1987). A ROWS is a ROSE: Spelling, sound, and reading. Memory & Cognition, 15(3), 181–198. 10.3758/BF03197716

57. Wagner, A. D., Paré-Blagoev, E. J., Clark, J., & Poldrack, R. A. (2001). Recovering meaning: Left prefrontal cortex guides controlled semantic retrieval. Neuron, 31(2), 329–338. 10.1016/s0896-6273(01)00359-2

58. Wing, E. A., Marsh, E. J., & Cabeza, R. (2013). Neural correlates of retrieval-based memory enhancement: An fMRI study of the testing effect. Neuropsychologia, Special Issue on Functional Neuroimaging of Episodic Memory, 51(12), 2360–2370. 10.1016/j.neuropsychologia.2013.04.004

59. Zhuang, L., Wang, J., Xiong, B., Bian, C., Hao, L., Bayley, P. J., & Qin, S. (2022). Rapid neural reorganization during retrieval practice predicts subsequent long-term retention and false memory. Nature Human Behaviour, 6(1), 134–145. 10.1038/s41562-021-01188-4

