## Supplementary material for "Memory Traces Reflect How They Were Last Accessed": Supplementary_Materials_revised.docx

**Text S1. Impact of Re-exposure Accuracy on Recognition Accuracy**

We examined whether final recognition accuracy depended on whether a pairing had been correctly accessed during re-exposure. For each participant, we categorized pairings based on whether the second re-exposure trial was answered correctly, and compared recognition accuracy between the two sets of pairings. Recognition accuracy did not differ between correctly and incorrectly accessed pairings (t(19) = 1.22, p = 0.237). Because corrective feedback was provided after every re-exposure trial, a pairing answered incorrectly during re-exposure was followed by a learning opportunity before recognition. Therefore, re-exposure accuracy might not be a good predictor of recognition performance, consistent with the rest of our behavioral findings**.**

**Text S2. Covariation in Classifier Confidence Across ROI**

As a follow-up to the ROI-based classification analysis, we examined whether regions that successfully decoded prior semantic access history showed coordinated trial-wise fluctuations in classifier confidence, also using an informational connectivity approach. This analysis was restricted to ROIs that achieved above-chance classification performance (VT, EVC, and hippocampus). For each participant and ROI, GNB classifiers were trained using only correctly recognized trials. For each test trial, we extracted the posterior probability assigned to the true class as the continuous confidence metric, and concatenated confidence values across folds to form one confidence vector per ROI and participant. Covariation in classifier confidence between pairs of ROIs was quantified as Spearman correlations between the confidence vectors, which were Fisher-Z transformed before group-level analysis. Statistical significance was calculated using a permutation test in which confidence values from one ROI were randomly shuffled within run while confidence values from the other ROI were held fixed. This procedure was repeated 10000 times to generate a null distribution of group-averaged covariation values.

Among the three ROIs that showed above-chance classification performance, classifier confidence showed significant trial-wise covariation across all pairs. Specifically, confidence vectors were strongly correlated between EVC and VT (mean rho = 0.51), and moderately correlated between EVC and hippocampus (mean rho = 0.29) and VT and hippocampus (mean rho = 0.32). All observed correlations exceeded the corresponding null distributions generated by permutation testing (p<0.001). These results indicate that regions supporting above-chance decoding of prior semantic access history also showed coordinated fluctuations in classifier confidence across trials.

**Table S1. Recognition Pattern Consistency t test results in all ROIs.** ** = p_FDR_ < 0.01, *** = p_FDR_ < 0.001

| **ROI** | **Semantic access condition** | **t** | **p_FDR_** |  |
| --- | --- | --- | --- | --- |
| VT | item | 0.31 | 0.79 |  |
| VT | category | -1.33 | 0.54 |  |
| VT | theme | 0.32 | 0.79 |  |
| EVC | item | -0.75 | 0.79 |  |
| EVC | category | 7.22 | <0.001 | *** |
| EVC | theme | -0.57 | 0.79 |  |
| ATL | item | -1.89 | 0.29 |  |
| ATL | category | -0.78 | 0.79 |  |
| ATL | theme | -0.57 | 0.79 |  |
| Hippocampus | item | -1.82 | 0.29 |  |
| Hippocampus | category | 0.33 | 0.79 |  |
| Hippocampus | theme | -1.89 | 0.29 |  |
| PrC | item | -2.24 | 0.23 |  |
| PrC | category | 0.53 | 0.79 |  |
| PrC | theme | -1.11 | 0.67 |  |
| vlPFC | item | -2.37 | 0.23 |  |
| vlPFC | category | -0.32 | 0.79 |  |
| vlPFC | theme | 0.58 | 0.79 |  |
| MPC | item | -0.46 | 0.79 |  |
| MPC | category | 0.06 | 0.95 |  |
| MPC | theme | -1.31 | 0.54 |  |
| VWFA | item | -4.11 | 0.01 | ** |
| VWFA | category | 0.37 | 0.79 |  |
| VWFA | theme | 0.41 | 0.79 |  |

**Figure S1 Recognition consistency results in all ROIs.**

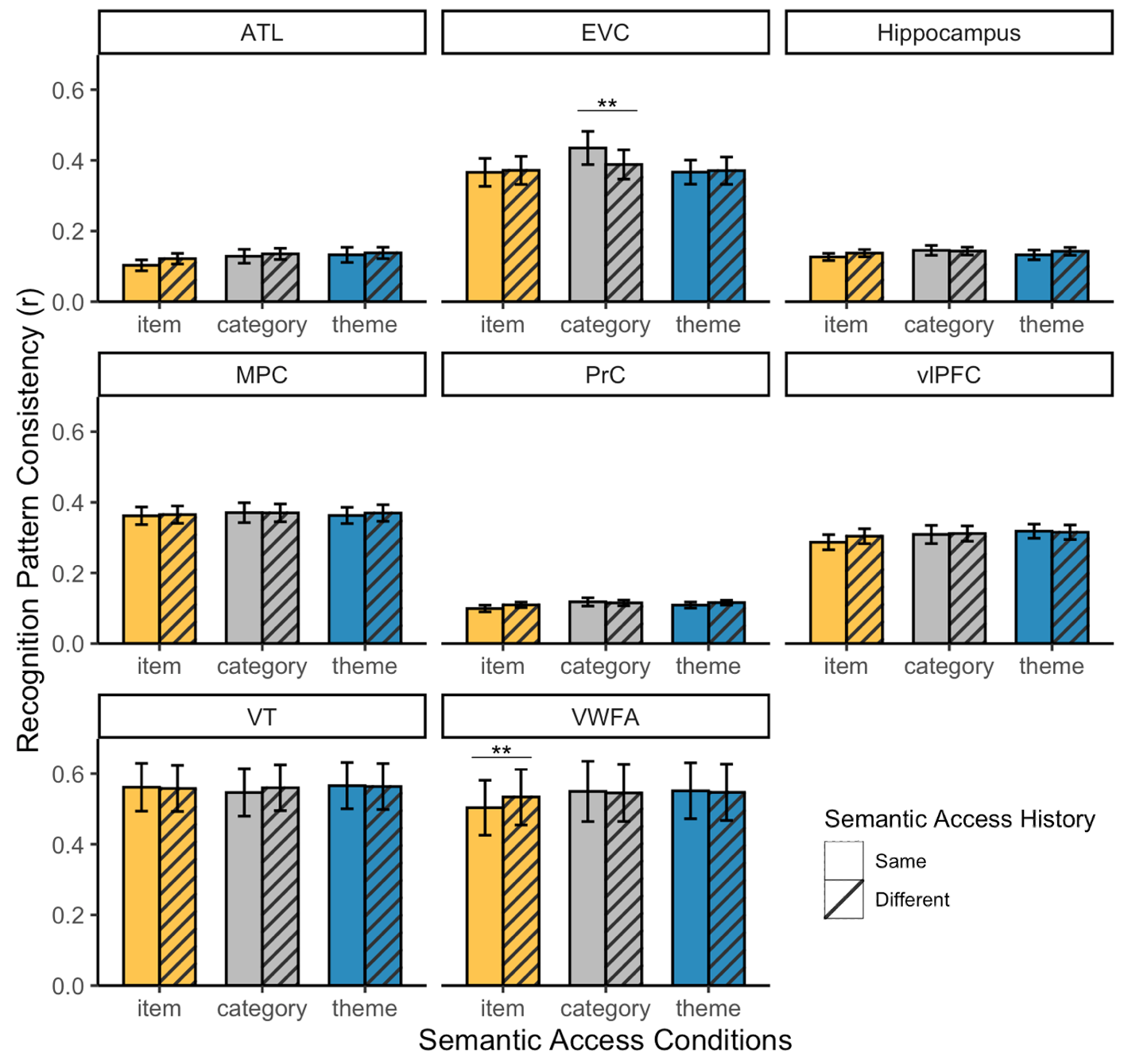
The recognition neural patterns for items previously accessed through the same (within) versus different (between) types of semantic access. Spearman’s correlation coefficients standardized with Fisher’s z. *p_FDR_ < .05, **p_FDR_ < .01, ***p_FDR_ < .001.
